# Long-term evaluation of the association between dominance, bib size, sex and age

**DOI:** 10.1101/2025.02.13.638048

**Authors:** Nicolas J. Silva, Liliana R. Silva, Pietro B. D’Amelio, André C. Ferreira, Annick Lucas, Rita Covas, Claire Doutrelant

**Author notes:** **Corresponding author:** Nicolas J. Silva.

## Abstract

The association between dominance and phenotypic traits (‘badges of status’) has been recently questioned, since it has been found to show more variation across populations and time than expected. Using an eleven-year dataset, encompassing more than 1,800 individuals, we studied the association between the size of a melanin-based plumage trait (a black patch under the beak or ‘bib’), individual attributes (age and sex) and dominance. Our study model, the sociable weaver (*Philetairus socius*), is a small passerine, which lives year-round in colonies of varying size. Bib sizes were obtained from pictures and automatically pre-processed with an object segmentation deep learning network. We recorded 36,346 dominance-related interactions to estimate dominance hierarchies. We then investigated the effect of sex and age on bib size, and the overall and between-year association of bib size and age with dominance for each sex. We found that bib size increased with age within individuals of both sexes. Age was a strong and consistent predictor of dominance in males, but not in females. Nevertheless, dominance was not correlated with bib size in males and in females. Finally, these relationships did not strongly vary between years. Our results suggest that males and females may use different factors to regulate dominance interactions. Furthermore, our longitudinal dataset allowed us to show that bib size is not a badge of status in our population even though its growth encodes information on age. These results highlight the importance of considering multiple traits over time when studying morphological signalling of dominance hierarchy.

## 1. Introduction

Competition over resources (e.g., food, territories, mates) often lead to aggressive interactions between individuals. These agonistic events can be very costly, sometimes leading to severe injuries or death (Clutton-Brock et al., 1979; Neat et al., 1996; Riechert, 1998; Sneddon et al., 2002). To limit these costs, many species establish dominance hierarchies (Tibbetts et al., 2022). Several traits or individual attributes, most commonly sex or age, have been found to be linked to the position of individuals in the dominance hierarchy (Tibbetts et al., 2022).

Dominance signalling can rely on intrinsic competitive abilities of individuals and can be displayed through ‘badges of status’. These traits can be visual, olfactory or acoustic and should correlate with the individuals’ dominance status (Rohwer, 1975, 1982; Senar, 2006; Smith & Harper 2003; Whiting et al., 2003). They allow to resolve conflicts without direct confrontation, because individuals can assess the competitive ability of others before interacting physically with them. These traits are supposed to be honest and several non-mutually exclusive mechanisms have been shown to be part of the evolutionary stability of the honesty of the signal, among which condition-dependence, social punishment, physiological costs, predation costs, or familiarity (Chaine et al., 2018; Searcy & Nowicki, 2010; Senar, 2006). The signalling function of badges of status has been demonstrated both by correlational and experimental studies in a wide range of taxa (e.g., mammals (Burgener et al., 2009; Clutton-Brock & Albon, 1978), birds (Senar, 2006), reptiles (Stapley & Whiting, 2006), and insects (Tibbetts & Lindsay, 2008)).

In birds, a meta-analysis using 47 studies encompassing 23 species found that plumage was linked to dominance across species and plumage types (e.g., carotenoids, melanic), suggesting that there is overall statistical support for the status signalling hypothesis across birds (Santos et al., 2011); however, heterogeneity was high, indicating low generality of that finding across studies. Regarding melanin-based plumage patches in birds, the textbook example has been the bib size of male house sparrows *Passer domesticus*, which was found to be correlated with the fighting abilities of individuals (Nakagawa et al., 2007). However, this example was challenged in a meta-analysis conducted on 19 studies (Sánchez-Tójar, Nakagawa et al., 2018), which showed that when including unpublished studies, the overall effect size of the correlation between dominance and bib size was not different from zero, and that the overall evidence for the hypothesis in house sparrows has been declining over time. This meta-analysis revealed a strong publication bias that calls for reconsideration of the bib size-based status signalling in this species, and for further investigation of within-species variation in the relationships between badges of status and dominance rank.

In addition to dominance signalling through phenotypic traits, dominance hierarchies have also been shown to be associated with individual attributes, most notably age. The hierarchies associated with individual attributes are called ‘convention-based’ hierarchies, and in these, age influences various phenotypic traits and regulates dominance interactions in many taxa (e.g., in birds (Sergio et al., 2009; Stahl et al., 2001), insects (Taylor et al., 2020), and mammals (Šárová et al., 2013)). For instance, in chimpanzees (*Pan troglodytes*), female social status is influenced by age, while males fight to gain status in the male hierarchy, which ultimately increases their reproductive success (Foerster et al., 2016).

The mechanisms regulating dominance may differ between the sexes because the benefits (and costs) of dominance are unlikely to be equivalent, and sex-biased dispersal may lead to stronger social bonds within one of the sexes. In birds, status traits have been studied mostly in males and less in females (Doutrelant et al., 2020), but some studies, with relatively small sample sizes, on mutually ornamented species found an association between dominance and a given ornament in females (e.g., Chaine et al., 2011; Crowhurst et al., 2012; Jawor et al., 2004; Murphy et al., 2014; Watt, 1986). Hence, the selective pressures affecting these ornaments and their signalling function may be sex-specific. Finally, studies have also found evidence for a sex-dependent relationship between dominance and age (Foerster et al., 2016). However, overall sex differences in the mechanism regulating dominance remains to be explored in depth.

Trait association can also vary in time and according to environmental conditions (Kvarnemo & Ahnesjo, 1996; Lucchesi et al., 2019), and dominance is no exception. Therefore, being able to disentangle the effect of confounding factors, such as sex, age, or phenotypic traits on dominance and the variations through time is crucial to understand the evolution and the stability of dominance in wild populations. In a recent review, Tibbetts et al. (2022) argued that long-term field research is needed to deduce the factors influencing the stability of dominance hierarchies. However, collecting such data represents a challenge as it requires continuous funding to cover both logistical and human resources (Cockburn, 2014). Another challenge is the processing of the large volumes of data collected, which is very time consuming and costly when extracting information from videos or images (Weinstein, 2018). Recent approaches based on deep learning methods (LeCun et al., 2015) can help with image processing, as they allow to conduct time-consuming and repetitive tasks much faster than humans (e.g., few milliseconds per images classification; Jocher et al., 2022) and sometimes outperforming the repeatability and performance of humans (Ditria et al., 2020). In ecology, deep learning methods are increasingly used for multiple tasks such as identification, or density estimations, among others (Christin et al. 2019). However, these methods are still rarely used for fine scale measurements of morphological traits, as it requires accurate detection and precise segmentation of the phenotypic traits of interest.

In the present study, we examine the relationship between a putative badge of status (a black plumage ‘bib’) and dominance, age, and sex in a highly social colonial and cooperative bird, the sociable weaver (*Philetairus socius*). To this end, we used a deep learning approach, based on object-segmentation, to automatically extract the bib size from approximately 13,300 pictures. We then calculated dominance from more than 36,000 dominance interactions collected over seven years. In this species, both male and female adults display a black melanin-based bib, with males only presenting very slightly larger bibs (+0.032 cm²; Acker et al., 2015). Sociable weavers establish steep dominance hierarchies within their colonies with males being dominant over females (Rat et al., 2015; Silva et al., 2018) and older males have been found to be dominant over younger males (Silva master thesis, 2017). Previous short-term studies on this species found conflicting results regarding the association between dominance and bib size, with either a positive association in male-male interactions only (over two years; Rat et al., 2015), or females with larger bibs being dominant when interactions were computed in both sexes (Silva master thesis, 2017).

Here, we first evaluated how bib size varies in relation to sex and age, and performed a longitudinal analysis to understand the within individual variation in bib size, using 11 years of data. Then, over seven years of data, we tested how dominance related with bib size, age and sex, and whether these associations varied between years. Since dominance hierarchies in this species are structured by sex (with most males being dominant over most females), we then explored whether the traits associated with dominance hierarchies were different when estimating all individual compared to when estimating same-sex dominance hierarchies. Longitudinal studies on badges of status are rare and we expected that by using such a large dataset spanning across time, we could reliably assess whether bib size functions as a badge of status, as well as capture potential variation over time of the relationship between dominance, age and bib size.

## 2. Materials and Methods

### 2.1. Species and individual information

Sociable weavers are passerines endemic to the semi-arid savannahs of the Kalahari region of southern Africa. They are small sexually monomorphic birds of approximately 28g and are colonial facultative cooperative breeders (Maclean, 1973). The study was conducted at Benfontein Nature Reserve, in South Africa (-28°818S, 24°816E, 1,190m asl), an area of ∼15km² of open savannah where up to 15 colonies are simultaneously monitored as part of a long-term study since 2010 (some colonies collapsed during the course of the study and new ones were added to the long-term monitoring).

The study was conducted between 2013 and 2024. Dominance trials were conducted on seven of the regularly monitored colonies (from 2013 to 2019), while bib pictures were taken from all colonies (from 2013 to 2024). We define a year as the period between 1^st^ of September and the 31^st^ of August, as the birds in the study colonies are captured at the end of the Southern Hemisphere winter, generally in August-September, and the breeding season usually starts in September-November (D’Amelio et al., 2024b). Sociable weavers can form very large colonies up to approximately 500 individuals (Maclean 1973), but colonies in our study area over the years of this study varied between 7 and 82 birds (mean±SD, 38±17 individuals). Individuals are ringed with three coloured rings and a numbered metal ring, allowing individual visual identification. We also take morphological measurements (mass, tarsus+intertarsal joint’s length) and blood samples to sex genetically all individuals (D’Amelio et al., 2024a; Griffiths et al., 1998).

Exact age is known for all individuals that were born in the studied colonies as a result of a routine breeding monitoring (these individuals constitute 77% of the males and 65% of the females in the dataset). For immigrants, which were first captured as adults, we computed the age by adding the “average minimum age” of first dispersal to the first date of identification. This average minimum age of first dispersal in days was quantified for both males and females based on the long-term capture-recapture database (mean±SD, males: 690±46 days, N=13; females: 727±291 days, N=184). In sociable weavers, females disperse away from their natal colonies to breed (Covas et al., 2006; van Dijk et al., 2015). Once they establish themselves in a colony, they generally settle for several years and no longer change colonies ((mean±SD) 2.9±1.6 of years of presence in the study site after moving in from the outside and starting to breed, N=562 females; Silva et al., in prep). Note that our estimation of age is more robust for females (N=184) than for males (N=13). All protocols were approved by the Northern Cape Nature Conservation (permit FAUNA 1638/2015, 0825/2016, 0212/2017, 0684/2019 and 0059/2021) and the Ethics Committee of the University of Cape Town (permit 2014/V1/RC and 2018/V20/RC).

### 2.2. Dominance status

Dominance hierarchies were determined within seven colonies each year (2013-2019) by analysing and scoring agonistic interactions between individuals feeding on an artificial food source, a plate with mixture of bird seed, placed beneath each colony. Behavioural observations were performed using a video camera (Sony Handycam HD) on a tripod 2-3m from the feeder, to record all interactions seen approximately up to 1m around the feeder. All encounters were quantified as either aggression, displacement, threat or avoidance (see Table 1 in Silva et al., 2018 and Rat et al. 2014), and for each we determined the winner and the looser of the interaction.

**Table 1:** Estimates of the model parameters to test the influence of the size and the age on the bib size using the whole dataset. Bib size was taken once a year at the end of winter and some individuals were collected multiples years. The model was computing using 3,831 observations (995 females and 868 males). Here, ‘Age’ is either exact age for individuals born on our studied colonies or minimum age for individuals first caught as adults. Conditional R² was of 0.67.

| Fixed effect | Estimate | SE | t-value | p-value |
| --- | --- | --- | --- | --- |
| **Intercept** | 1.55 | 0.03 | 45.91 | <0.001 |
| **Age** | 0.05 | 0.00 | 9.78 | <0.001 |
| **Sex - Male** | 0.02 | 0.01 | 2.58 | 0.010 |
| **Tarsus** | 0.02 | 0.00 | 4.48 | <0.001 |
| **Age : Sex - Male** | -0.01 | 0.01 | -0.86 | 0.392 |
| Random effect | Variance |  |  |  |
| **Individual ID - Intercepts**<br>(N=1,863) | 0.013 |  |  |  |
| **Individual ID - Slopes** (N=1,863) | 0.001 |  |  |  |
| **Colony** (N=19) | 0.001 |  |  |  |
| **Year** (N=10) | 0.002 |  |  |  |
| **Photographer** (N=7) | 0.004 |  |  |  |
| **Camera** (N=3) | 0.001 |  |  |  |
| **Residuals** | 0.012 |  |  |  |

Dominance observations were performed most of the time in September at the end of winter, before the onset of the breeding season (see Table S15 for precise dates). From 2014 to 2017 and in 2019 observations were conducted in September; in 2013, from September to November; and in 2018, in December. Over these seven years, we conducted approximately 4h of observations each day, usually starting between 06:45 and 12:50; and repeated later in the day, between 13:15 and 16:50, and the food provided was removed between recordings. We obtained a total of 550 hours of video and each colony was recorded approximately (mean±SD) 15±12 hours per year (mean±SD, over approximately 6±4 different days per colony). All the videos per colony per year were recorded within a period length of (mean±SD) 9±9 days per year (Table S15). A total of 36,346 agonistic interactions (6,869 acts of aggression, 14,856 displacements, 3,459 threats and 11,162 avoidances) were recorded (mean±SD, approximately 1,010±1,240 interactions per colony per season). In addition, we scored 682 agonistic interactions for which we were not able to determine the winner or loser, as both individuals, after initiating a strength assessment, move away simultaneously from the confrontation.

To assess individual rank and to obtain dominance hierarchies, we used David’s score (David, 1987), which is considered as one of the most appropriate and reliable methods available (Sánchez-Tójar, Schroeder et al., 2018), as it allows to assess overall individual success in agonistic interactions (Gammell et al., 2003). David’s score allows to estimate dominance of an individual by considering the proportion of wins or losses across all the agonistic interactions. This index is also adjusted by the total number of observations of the individuals and the proportion of wins for each of them based on the relative number of wins and losses against all the other individuals in the group. Additionally, this method allows to compare results to previous studies on dominance behaviour in this species (Rat et al., 2015; Silva master thesis, 2017). This score is calculated using the “steepness” R package (De Vries et al., 2006) and standardized per year and colony between zero and one to allow comparison between individuals of different groups (see Fig. S1 for the distribution of the dominance scores in the population).

The hierarchies produced here included the majority of the birds captured in winter for the studied colonies ((mean±SD) 85%±23% of individual within colony and year observed during dominance data collection; Table S15). The 15% missing correspond to birds never observed interacting at the feeders or that were not alive/present at the colony at the recording time. Individuals seen in several colonies (N=29 individuals) and those seen interacting less than three times in a given colony and year (N=45 individuals) were excluded from the analyses to avoid prospecting individuals (i.e., individuals briefly visiting colonies without being part of it).

In addition to these general dominance hierarchies (both-sexes dominance hierarchies) including all individuals and interactions, we also extracted dominance hierarchies by considering i) only male-male dominance interactions (N=16,261 interactions between 256 males, the between-males hierarchies) and ii) only female-female dominance interactions (N=2,651 interactions between 267 females, the between-females hierarchies).

To obtain reliable and representative dominance hierarchies for steep dominance hierarchies, like the sociable weavers’ (Silva et al., 2018), Sánchez-Tójar, Schroeder et al. (2018), proposed that the threshold ratio of total number of observed interactions to total number of individuals should have a minimum of 10. In our study site, many colonies did not fulfil this criterium. We therefore decided to run our models both with and without the colonies with a ratio inferior to 10 (except for females as only one colony fulfilled this criterium). We found that the estimates were similar between the models with and without the colonies with a ratio inferior to 10 (Table S3 to S12), and therefore decided to include the colonies with a ratio inferior to 10 in the results.

### 2.3. Bib size and YOLO: annotations, training bib segmentation and model validation

For all the birds captured, three top view pictures of the bib per individual were taken in the field (Rat et al., 2015). We collected pictures of the bib of the birds for 11 years, between 2013 and 2024, between mid-August and mid-September each year. The birds were handled and held in an apparatus where they were photographed in a standardized position. Pictures of the bib and videos for dominance estimation of the birds were collected with less than a month from each other for all years, except for three colonies of the year 2013 (approximately 1.5 month between data collection) and four colonies of the year 2018 (approximately 3.5 months between data collection).

In previous works, bib size was manually analysed using image graphic editor (The GIMP Development Team, 2019 or Photoshop CS5.1 (Version 12.1), hereafter referred as “GIMP method”), which is a time-consuming task when dealing with large datasets. We improved this method by implementing a deep learning approach using YOLO (You Only Look Once; Redmon et al., 2016) to automatically detect and extract bib size from the pictures (Supplementary 2 for detailed description of the method).

We tested the repeatability of the GIMP and YOLO methods by selecting 20 individuals (300 pictures) on five different years for which bib size was measured with both methods (rptR package version 0.9.22 (Nakagawa & Schielzeth, 2010)). Results showed high repeatability and high correlations between the two methods (Table S1), indicating that the bib size measurements obtained with the deep learning model were robust. Bib size, extracted in pixels from pictures, was transformed to cm².

### 2.4. Statistical analysis

All statistical analyses were conducted in R version 4.3.1, using mixed models ran with the lme4 package version 1.1-33 (Bates et al., 2014). We verified whether model assumptions were met, specifically that a) the residuals followed normality using Kolmogorov-Smirnov tests, b) the simulated dispersion of the residuals was equal to the observed dispersion, c) there were no more simulated outliers than expected, and d) there was homoscedasticity of the variance at three quantiles (0.25, 0.5, 0.75) between the residuals and predicted values. These diagnostics were performed using the Dharma package version 0.4.6 (Hartig & Lohse, 2022), and, although not marked, some heteroscedasticity was observed and it is detailed in Supplementary 7 and 8. The plots were created with ggplot2 package version 3.4.2 (Wickham, 2014) and ggeffects version 1.3.2 (Lüdecke, 2023).

#### 2.4.1. Factors associated with bib size

To design and interpret the most appropriate structure for the models testing the association between bib size and dominance, we first needed to understand the relationship of bib size with age within sexes (Fig. S2). These variables were found to be associated in a previous study conducted on this species (using distinct datasets: Acker et al., 2015). In addition, we included tarsus length (as a proxy of body size), to correct for a potential correlative effect between bib size and body size (as recommended by Wysocki et al., 2022). We designed a linear mixed model (LMM) with bib size (in cm²) as the response variable, and tarsus length (in mm), age (in days), sex and the interaction between age and sex as fixed factors. Age is often used as a non-linear factor in many model to test for a senescence effect on the association between age and a trait (here bib size). However, here we did not test for such effect due to the low number of old individuals in our dataset. The oldest birds alive recorded on our study site was approximately 16 years old (D’Amelio et al., 2024a), but here our individual are on average three years old ((mean±sd): males - 1,212±865 days old and 20 individuals older than 10 years only; females - 1,057±739 days old and 10 individuals older than 10 years only). Therefore, we decided that including age, as a non-linear factor, would not be relevant and would not allow to test for floor or ceiling effect. As random factors, we controlled for year, colony and individual identity. Because photographers did not always follow the same protocol and because we changed cameras over the years, we included photographer ID in our initial model, the observer who took the photos (7 observers) and the camera model used (3 cameras). We also added random slopes to the random intercepts to estimate the link between age and bib size for every individual. We estimated random slopes uncorrelated with random intercepts, as we do not have predictions about individuals with larger bib size also having a greater bib size growth when getting older. Using the estimated slopes for each individual, we assessed the percentage of individuals showing positive growth in bib size over the years. Additionally, to evaluate the proportion of variance explained by the random intercepts and random slopes in the model, we computed two alternative models: (i) one excluding the random slopes and (ii) one excluding both the random intercepts and random slopes. The conditional R² (i.e., the proportion of variance explained by both the fixed and random effects) was calculated to estimate the proportion of variance explained by the random slopes and random intercepts when removed from the models (Johnson, 2014).

Tarsus length was computed as the average tarsus length measured during the adult life of the individuals (by the same scorer, R.C., which measured the majority of birds, or by others if no measures by R.C. were found for that individual). Three individuals with extreme tarsus length observations were removed.

We used a dataset composed of 1,863 individuals from 19 colonies over eleven years (from 2014 to 2024, N=3,831 observations). We estimated the influence of the number of repeated measures per individuals on the random slopes variance estimation by computing the same model on a subset of individuals with at least three measures through their life and we obtained similar conclusions regarding the random slopes analysis (Table S3). In addition, we used 1,000 bootstraps and 1,000 permutations and estimated the between individual adjusted repeatability of bib size across years, using the ‘rptR’ package (Stoffel et al., 2017), on 959 individuals seen at least twice over the study period (2,927 observations). To estimate the repeatability, we only corrected for the sex of the individuals.

The residuals of the model using the 3,831 observations did not respect the normality assumption (Kolmogorov–Smirnov test, P<0.001). This non-normality of the residuals of the model were mainly due to one year photographed by one scorer. By removing this scorer from the dataset (N=647 observations), the model fitted the necessary assumptions. Estimates and effects of the different factors ranged from highly similar to identical between the two models (Table 1 and Table S2). Therefore, we decided to use the model containing the full dataset.

#### 2.4.2. Overall association between dominance, sex, bib size and age

First, we used 3,000 bootstraps and 3,000 permutations and estimated the between individual adjusted repeatability of dominance rank within the colonies across years, using the ‘rptR’ package (Stoffel et al., 2017), on 184 individuals observed at least twice over the study period (498 observations). To estimate the repeatability, we corrected for the sex of the individuals and used a Poisson log-link model.

Then, we performed a LMM using the dominance score of the individuals as the response variable and, as fixed factors, bib size, age, sex, tarsus length, body index, as well as the interactions between bib size and sex, and age and sex. Gaussian distribution model was used as they fitted particularly well (see Supplementray 6 Fig. S4). Body index was based on a scaled mass index (Peig & Green, 2009) that uses mass (here collected at the capture events, approximately when dominance data collection was performed) and a body size metric of an individual (here tarsus length, T) with its arithmetic mean value for the studied population (T_0_). The method allows to compute a scaled mass as the predicted body mass when T=T_0_. To visually describe the structure of our model, we designed a Directed Acyclic Graph (DAG) (Fig. S2). DAGs are visualisation tools that help to represent the links between the variables that we intend to test in a model (Wysocki et al., 2022). In this model, bib size was normalized within colony and year when analysing both-sexes dominance hierarchies, and within colony, year and sex when dealing with sex-specific dominance hierarchies. The normalisation allows us to compare relative bib size of the individuals within colonies and does not allow us to interpret bib size at the population level. Two individuals with a David score of one in different colonies might not have the same bib size but potentially be at the same bib size position in the bib size hierarchy within their colonies. Therefore, they would be assigned the same normalized bib size. Normalisation was performed as 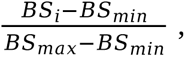 where *i* represents the individual’s bib size within the colony, and BS*_min_* and BS*_max_* respectively the minimum and maximum bib size in cm^2^ within a colony one year. Normalized bib size, body index, tarsus length and age were scaled and mean-centered over the whole dataset. Body index and tarsus length were included because they are often related to dominance in other bird species (Jonart et al., 2007; Tibbetts et al., 2022), and including them allowed to estimate the influence of bib size, while accounting for the influence of the individual’s condition and its size on dominance score (Fig. S2 for visual representation of the links between dominance and the predictors). As random factors, we controlled for colony (N=7), individual identity (N=409) and year of data collection (N=7, from 2013 to 2019). We included year as a random factor since we expected potential differences in dominance between years (e.g., arising from differences in social composition, resource availability, breeding onset).

#### 2.4.3. Between-years association between dominance, sex, bib size and age

We used the same LMM model structure as when testing the overall association between dominance and bib size, but instead of having “year” as a random categorical factor, we used it as a fixed categorical factor. Additionally, we included the interactions between year, normalized bib size and sex, as well as between year, age and sex. We then performed a post-hoc test with the emmeans package version 1.8.6 (Lenth et al., 2023) to determine if the slopes of the relationship between dominance, bib size and age were significantly different from zero between years (Table S13).

## 3. Results

### 3.1. Factors associated with bib size

We found that age and bib size were significantly positively associated (0.02±0.01 (EST±SE, used throughout the Results section except if specified otherwise), T=2.58, P=0.010; Table 1), and we estimated that between individuals that differ by one year, the older birds have approximately 2% larger bibs. Within individuals, we found a positive association between bib size and age for 99.8% of the birds (Fig. 1), and this relationship was not affected by sex, with bib size increasing by 1.37% per year within females and 1.21% per year within males. We found a conditional R² of 0.67 in the model including both random intercepts and random slopes, and estimated that the individual random intercepts for bib size accounted for more variance in the model (Conditional R² = 0.64 in the model without random slopes) than the random slopes of age’s effect on bib size per individual (Conditional R² = 0.26 in the model without random intercepts and slopes). This suggesting that individuals vary in their average bib size but have a similar pattern of growth through time. Moreover, we found a small positive significant effect of sex (0.02±0.01, T=2.58, P=0.010; Table 1) with males having on average 1.29% larger bib size than females (mean±SD, 1.55±0.03 cm² for females and 1.57±0.03 cm² for males). Finally, we found a significant positive association between tarsus length and bib size (0.02±0.00, T=3.48, P<0.001; Fig. S3; Table 1), hence larger individuals have larger bibs independently of sex. Finally, bib size was found repeatable between individuals and across years ((estimate [CI]) 0.41 [0.37; 0.44], P = 0.001).

**Figure 1:**
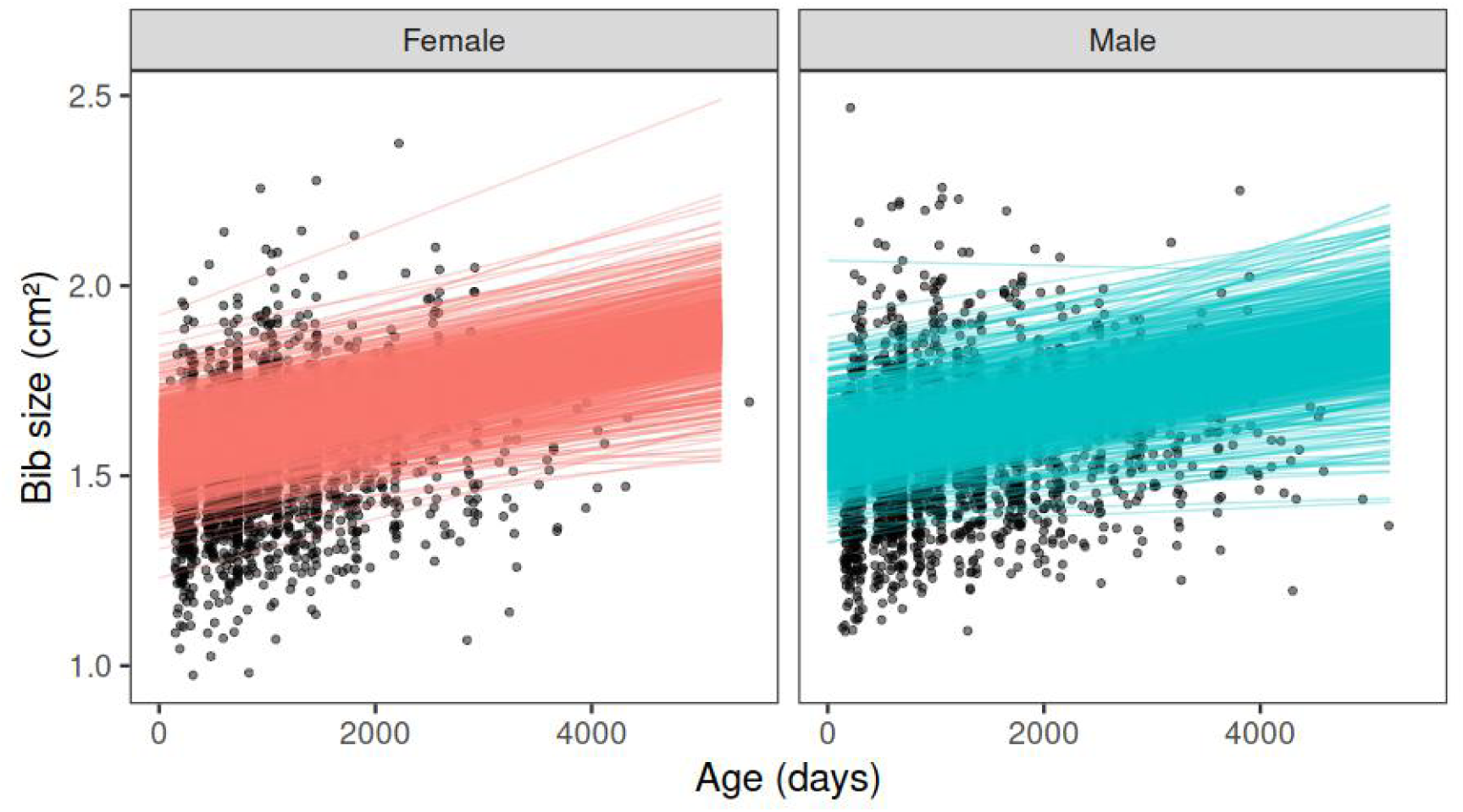
Linear regression of bib size as a function of age. The random slopes and intercepts of the 1,863 individuals (995 females (red) and 868 males (blue)) were used to simulate bib size development every 200 days between one to 5,201 days old (i.e., more than 14 years).

### 3.2. Overall association between dominance, bib size, age and sex

We found a between individual repeatability of dominance rank of 0.38 ([CI] [0.28; 0.49], P=0.001).

#### Dominance in relation to bib size and sex

We found no significant association between normalized bib size and dominance in the datasets considering both-sexes dominance hierarchies (0±0.01, T=0.19, P=0.852; Table 2; Fig. 2.A) and between-males dominance hierarchies (-0.02±0.01, T=-1.69, P=0.092; Table S5; Fig. 2.C). However, we found a small positive significant correlation between bib size and dominance in between-females dominance hierarchies (0.04±0.02, T=2.34, P=0.020; Table S7; Fig. 2.E). Females being ranked 25% higher in the normalized bib size ranks have an estimated 8% higher dominance score. However, the between-years analysis below shows that the relationship is mainly influenced by two years with very low sample size (Table S13) and when removing these two years the correlation is weak and non-significant (0.02±0.02, T=1.06, P=0.290, Table S14).

**Figure 2:**
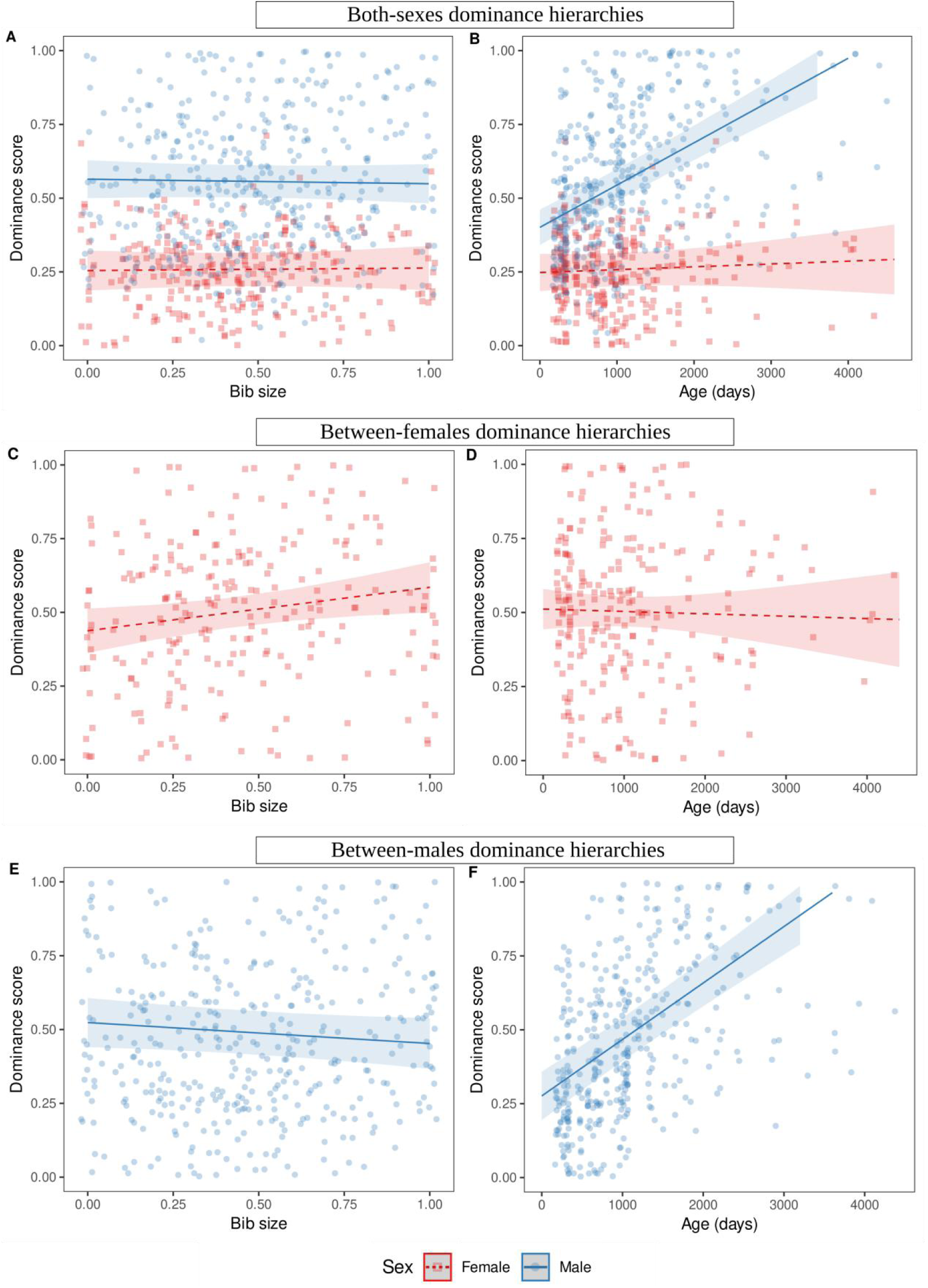
Association between: (A,C,E) dominance and normalized bib size; and (B,D,F) dominance and age. Results were obtained using both-sexes dominance hierarchies (A,B), between-females dominance hierarchies (C,D) and between-males dominance hierarchies (E,F). Dominance score and bib size were normalized within dominance hierarchies (both-sexes, between-females, and between-males), colony and year of data collection. Females are represented in red (square dots and dashed line) and males are represented in blue (circle dots and solid line).

**Table 2:** Estimates of the model testing the overall influence of bib size, sex and age on dominance in both-sexes dominance hierarchies, with colonies with a ratio inferior to 10 of interactions to the number of individuals. The model was computed with 723 observations. Association of dominance with age, bib size and sex is visually represented in. **Fig. 2.A and 2.B**.

| Fixed effect | Estimate | SE | t-value | p-value |
| --- | --- | --- | --- | --- |
| **Intercept** | 0.26 | 0.03 | 9.00 | <0.001 |
| **Bib size** | 0.00 | 0.01 | 0.19 | 0.852 |
| **Sex - Male** | 0.30 | 0.02 | 18.61 | <0.001 |
| **Age** | 0.01 | 0.01 | 0.65 | 0.515 |
| **Body condition** | 0.02 | 0.01 | 1.89 | 0.059 |
| **Tarsus** | 0.01 | 0.01 | 1.13 | 0.259 |
| **Bib size : Sex - Male** | -0.01 | 0.01 | -0.42 | 0.674 |
| **Age : Sex - Male** | 0.11 | 0.02 | 7.20 | <0.001 |
| Random effect | Variance |  |  |  |
| **Individual ID (N=409)** | 0.010 |  |  |  |
| **Year (N=7)** | 0.002 |  |  |  |
| **Colony (N=7)** | 0.003 |  |  |  |
| **Residuals** | 0.022 |  |  |  |

#### Dominance in relation to age and sex

In both-sexes dominance hierarchies, we found a strong significant link between sex and dominance, indicating that males are approximately two times more dominant than females (0.30±0.02, T=18.61, P<0.001; Table 2). We also found a strong association between dominance and the interaction of age and sex (0.11±0.02, T=7.2, P<0.001; Table 2), indicating a different relationship between age and dominance for males and females (Fig. 2.B), specifically a much steeper slope for males. Dominance score is 1.36% higher in females and 9.39% higher in males when comparing two different individuals that differ by one year in age (Table 2). This was found also in the sex-specific interactions’ datasets, where we observed a different association between age and dominance, with a positive, strong and significant link in between-males dominance hierarchies (0.16±0.01, T=11.77, P<0.001; Table S5; Fig. 2.D), while there was no statistically significant relation in between-females dominance hierarchies (-0.01±0.02, T=-0.35, P=0.725; Table S7; Fig. 2.F).

### 3.3. Between-years association between dominance, bib size, age and sex

#### Dominance in relation to bib size, sex and year

We found no evidence that the relationship between dominance and normalized bib size varied between years, with a few exceptions. Out of the seven years used in each analysis, the only statistically significant associations we found were a slightly negative relationship for one year in males in both-sexes dominance hierarchies (-0.06±0.03, T=-2.18, P=0.030; Table S13; Fig. S5.A in year 2016), a slightly negative relationship for one year in between-males dominance hierarchies (-0.06±0.03, T=-2.04, P=0.042 in year 2014; Table S13; Fig. S5.E), and positive relationships for two years in between-females dominance hierarchies (0.14±0.06, T=2.34, P=0.020 in year 2013, and 0.17±0.08, T=2.25, P=0.026 in year 2016; Table S13; Fig. S5.C). However, the sample size was limited for these latter years, with respectively 19 and 12 individuals only (the other years presenting between 17 and 84 individuals, Table S13), and so these relationships remain unclear.

#### Dominance in relation to age, sex and year

We found no between-years variation in the association between dominance and age in both-sexes dominance hierarchies and in between-males dominance hierarchies, with dominance being positively associated with age in all years for males and never associated for females (Table S13; Fig. S5.B and S5.F). In between-females dominance hierarchies, dominance was negatively associated with age in one year only (-0.07±0.03, T=-2.15, P=0.033; Table S13; Fig. S5.D in year 2017).

## 4. Discussion

We examined whether bib size and age relate to dominance at artificial feeders in sociable weavers and how these relationships vary between sexes and years. We found that, in both sexes, bib grows with age, and that larger birds have larger bibs. We also confirmed that males were dominant over females. However, normalized bib size was not found to be associated with dominance in any of the hierarchies (except in between-females dominance hierarchies for two years, but based on very low sample size). In addition, we confirmed that age was strongly associated with dominance in males a pattern that was consistent between years. In contrast, age was not found to be associated with female dominance. Our study is one of the largest long-term studies investigating correlates of dominance with a potential badge of status, providing robust results to understand the association between dominance and bib size, and the variation in its associated dominance correlates, given the low uncertainty in the variation obtained between years (see also Parker (2013); Sánchez-Tójar, Nakagawa et al. (2018); Tibbetts et al. (2022)). While previous studies found positive associations between dominance and bib size with only one year (included in this study (season 2013/2014); Silva master thesis, 2017) and two years of data (not included in this study; Rat et al. 2015), our analyses, on a much larger dataset, led to different conclusions. Finally, this study showed a sexual difference in the factor used to regulate dominance interactions.

### 4.1. Association between dominance, bib size and sex

Relative bib size was clearly not associated with dominance, whether when considering all birds or only males, and this did not change between years. We found an association within females, but the association was weak and unclear, with small effect sizes, and we had a smaller number of female-female interactions in the hierarchies (32 out of 33 colonies were below the recommended threshold of ratio of 10 of interactions to number of individuals in the group; Sánchez-Tójar, Schroeder et al. 2018). In addition, this association was only present in two years, which had among the lowest sample sizes of all years tested here (respectively N=19 and N=12 females). When these two years are removed there is no longer evidence of an association between relative bib size and dominance in females.

Several factors might explain why bib size does not function as a badge of status in this population. First, the sampling period could influence our results, as the association between bib size, age and dominance could be used by birds only in some parts of the season or some seasons depending on the social and environmental contexts. For example, a density-dependent effect in the association between badge of status and dominance has been found in the black-crested titmouse (*Baeolophus atricristatus*), where males with larger black crests had more access to feeding stations when competition was high but did not display such advantages when population density decreased and competition was low (Queller & Murphy, 2017). Here, the dominance scores were always computed at the end of winter, usually before the onset of the breeding season. Therefore, the association between dominance and bib size should be investigated at another moment of the year to investigate the possibility that bib size is a seasonal badge of status. Second, here we only tested the size of the bib, but other bib features (e.g., saturation, intensity) could signal the dominance abilities (Santos et al., 2011). For instance, in black-capped chickadees (*Poecile atricapilla*), high-ranking males exhibit darker plumages than subordinates (Poston et al., 2005). Finally, when facing known opponents, animals are expected to use information on the opponents’ previous behaviour (i.e., their ‘reputation’) to decide how to interact (Senar, 2006; Tibbetts & Dale 2007). For instance, Chaine et al. (2018) experimentally manipulated plumage patch size of golden-crowned sparrows (*Zonotrichia atricapilla*) and showed that manipulation only affected interactions between unknown individuals. Sociable weaver groups live together all year long, interacting on a daily basis, and this familiarity between individuals could remove the need for badges of status. In addition, male sociable weavers are philopatric, therefore knowing each other since birth. A previous study showed that the number of aggressive interactions between two individuals was negatively correlated with relatedness (Rat et al., 2015). Therefore, the social system of sociable weavers might not promote the use of badges of status in males. However, badges of status might still be important for individuals that move between colonies, especially females, which disperse to reproduce in other colonies (Covas et al., 2006; van Dijk et al., 2015). Badges of status could also play a more important role among the residents of very large colonies where individuals might not be able to keep track of all residents. Specifically, some sociable weaver colonies house several hundreds of birds (Maclean, 1973), while our study was conducted in colonies that varied in size between 7 and 82 residents.

Our results also contrast with previous results found in the same species but on smaller datasets, in which positive association between dominance and bib size was found in males (Rat et al., 2015) and in females (Silva master thesis, 2017). Accumulating evidence shows that phenotypic traits and their signalling function may vary depending on external forces (e.g., availability of food, mates), which vary over time and can ultimately influence competition intensity (Kvarnemo & Ahnesjo, 1996; Lucchesi et al., 2019). Long-term studies integrate the wide ecological and social variation that characterise the environments experienced by individuals (Clutton-Brock & Sheldon, 2010a, 2010b; Cockburn, 2014) and have provided several examples where limited temporal sampling could have resulted in inaccurate generalisation of association patterns (Clutton-Brock & Sheldon, 2010a; Cockburn, 2014; Cornwallis & Uller, 2010; Doutrelant et al., 2020). This study is an additional example illustrating the power of long-term studies in drawing general patterns of traits association.

### 4.2. Association between dominance, age and sex

We found a strong sexual difference in the association between age and dominance. Evidence of age-based dominance hierarchies (also referred to as social queuing) is common in many taxa and is particularly common in species that form long-term associations with relatives (Tibbetts et al., 2022), like sociable weavers. In ungulates, for instance, individuals, that are young and more inexperienced learn to be subordinates to older and stronger individuals, and do not challenge the hierarchy when getting older, resulting in age-based hierarchies (Favre et al., 2008; Thouless & Guiness, 1986). In sociable weavers, males know each other since birth and a mechanism similar to the ungulates could explain their age-based hierarchy. In contrast, females disperse to reproduce and therefore may not be able to use previous information about the status of dominant or subordinate of their opponents. Females may resort to alternative cues to infer fighting abilities of other females to decide how to interact with them, or may not need to enter in competition with them. It aligns with the low number of female-female interactions observed in this study, that is particularly low compared to the sex ratio observed in the population (Doutrelant et al., 2004). However, it is important to note that because of this natal dispersal of the females, our estimation of their age is less accurate than for males.

Social queuing has been suggested to also arise when the costs of reaching a higher position in the hierarchy are higher than the benefits of reaching that position (Broom et al., 2009). For instance, in chimpanzees, dominant males benefit from higher mating and reproductive success, therefore males fight for higher ranks (Foerster et al., 2016). However, females do not obtain fitness benefits from higher dominance rank in the hierarchy and intense fights would only bring fitness costs, resulting in age-dominance hierarchies in females (Foerster et al., 2016). In sociable weavers, males of the same colony are more related to each other than females and, within males’ fights could either bring direct fitness costs (if the fight is lost), but also indirect fitness costs (if the fight is won against a related individual). This could suggest that, for males, the fitness costs of reaching a higher dominance rank might not be compensated by the fitness benefits of reaching this rank, leading to age-based dominance hierarchies within males. In male sociable weavers, dominant individuals are more likely to access food and reproduce (Rat, 2015; Silva et al., 2018), but further information of the exact fitness costs and benefits on the long-term are needed to explore the origin of the age-based dominance hierarchies in males.

### 4.3. Association between bib size, age and sex

In agreement with a previous study (Acker et al. 2015), we found that bib size increases with age in both sexes. Age-related changes in adult plumage have been described in several species (e.g., Dreiss & Roulin, 2010; Wiebe & Vitousek, 2015). However, the bib size growth between years is small, only slightly over 1% for both sexes, and the perception by the individuals of such difference is questionable. Interestingly, we found that the growth rate of the bib is very similar between individuals, but individuals vary in bib size, indicating that the mechanisms behind bib size first growth varied between individuals. This last result is in line with the strong determination that genetics and development are expected to have on bird melanic patches (Roulin & Ducrest, 2013). Finally, we detected an effect of sex, with a difference of approximately +0.03cm² (approximately +2%) for the bib size of males compared to females. This sexual difference is in line with the one found previously in this species (Acker et al., 2015), but again, the perception of this difference by the birds would need to be demonstrated.

### 4.4. The growing importance of artificial intelligence in ecology

Finally, our methodology for measuring automatically thousands of bibs from pictures adds to the growing examples that illustrate the effectiveness and accuracy of deep learning-based methods to detect and quantify morphometrical traits, and how this can facilitate the process of extracting large amounts of data from images (i.e., here the mode processed 124 times faster than a human’s; based on 30 pictures). This approach can be used in different taxa, from museum specimens (Miele et al., 2020), to live ones (He et al., 2023), but it is still rarely used for such tasks in evolutionary ecology. While in many cases developing deep learning methods would be too demanding technically or too time consuming, data extraction from existing long-term datasets, as was the case here, is particularly well suited to be processed automatically (e.g., because of already laballed large amount of data).

## 5. Conclusion

In this long-term study capitalising on machine learning to assess the badge of status hypothesis in sociable weavers we found no convincing evidence of it, although for female-female interactions further investigation is needed. These results align with the previous recommendation of re-evaluating the status signalling hypothesis of melanic patches as well as evaluating the temporal variation of the correlates with dominance. By using multiple years of data, we were able to reach strong conclusions regarding the stability of bib size and age with dominance in our species. The sexual differences in the association between dominance and age for the two sexes suggest that male and female dominance interactions are not regulated by the same factors. These results show the importance of conducting analyses on both sexes when trying to understand the role of phenotypic traits on dominance, and suggest that males and females may be subjected to different selective pressures.

## Acknowledgments

We thank all involved in the long-term data collection and pre-processing over the years, specifically for help with the yearly captures, dominance videos and bib size picture collection. We thank Franck Theron for his management of the sociable weaver long-term database. We thank Margaux Rat, Inês Duarte and Bruna Fonseca for help scoring dominance videos. Additionally, we thank Margaux Rat for the initial development of the dominance and bib size collection and annotation protocols and Samuel Perret for bib size pictures and scoring. De Beers Mining Corporation provided access to Benfontein Nature Reserve and logistical assistance. The sexing and genotyping analyses were conducted at the CTM lab, CIBIO, and we thank Susana Lopes and all the technical staff that conduct the work. This study was funded by ANR to C.D. (France, grants 15-CE32-0012-02 and 19-CE02-0014-01), FCT (Portugal grants IF/01411/2014/CP1256/CT0007 and PTDC/BIA-EVF/5249/2014) ERC to R.C. (EU, Consolidator grant 866489) and by the DST-NRF Centre of Excellence at the Fitzpatrick Institute of African Ornithology University of Cape Town. N.J.S. was supported by the ANR 19-CE02-0014-01. L.R.S. was funded by ERC 866489 and ANR 19-CE02-0014-01. P.B.D.A. was supported by the south African Claude Leon Postdoctoral Fellowship 2019-20 and by the Marie Curie Individual Fellowship “SOCIAL MATCH” N. 896475 and then by the ANR 19CE02-0014-01. A.C.F was funded by University of Zurich Forschungskredit postdoc grant (K-74312-01-01 University of Zurich), Swiss Federal Commission for Scholarships and by ERC grants 866489 to R.C and. 850859 awarded to Damien Farine). R.C. was funded by FCT (IF/01411/2014/CP1256/CT0007 and CEECIND/03451/2018). The sociable weaver project is supported by the French OSU OREME, European Union’s Horizon 2020 Research and Innovation Programme under the Grant Agreement Number 857251, and the CNRS programme on Long-Term Studies in Ecology and Evolution (SEE-Life).

## Authors’ contribution

The study was initiated by C.D. and R.C., and N.J.S., L.R.S., P.B.D.A. and A.C.F. further contributed to its conceptualisation and execution. All authors contributed to the general monitoring data. L.R.S. and A.C.F. collected and processed the dominance score of the birds. N.J.S., L.R.S., A.L., R.C., and C.D. collected and processed bib data. N.J.S. developed the deep learning model. N.J.S. and P.B.D.A. produced the statistical analyses. N.J.S. led the writing of the first draft of the manuscript with contribution from all co-authors, and all authors contributed critically to the drafts and gave final approval for publication. P.B.D.A., R.C. and C.D. supervised N.J.S. for this work. R.C. and C.D. administrated the research project and acquired the funding for this work.

## Data availability statement

Analyses reported in this article can be reproduced using the data provided by Silva et al. (2025).

## Supplementary materials

This supplementary materials include:

i. Distribution of the normalised dominance score in the dataset (Fig. S1)
ii. The description and evaluation of the deep learning method used for automatic bib segmentation on pictures of the birds (Table S1).
iii. The evaluation of the relationship between bib size, age, and sex. Two models are presented: one with all the individuals; and one with a subsample due to non-validation of the mixed model assumptions in the first model (Table S2).
iv. The Directed Acyclic Graph of the model testing the association of dominance with bib size and age (Fig. S2).
v. The association between bib size and tarsus length (Fig. S3).
vi. Justification of the use of Gaussian distribution models to test for the association between dominance and bib size and age (Fig. S4).
vii. The summary of the mixed models testing the overall association of dominance with bib size and age in relation to sex. The models were computed using both-sexes dominance hierarchies (Table S3 and S4), between-males dominance hierarchies (Table S5 and S6) and between females dominance hierarchies (Table S7). For each dominance hierarchies, the models were computed both with and without the recommended threshold of ratio of interactions to number of individuals in the group superior to 10 (Sánchez-Tójar et al., 2018) (Table S4 and S6), except for female dominance hierarchies for which not enough colonies matched the criterion.
viii. The summary of the mixed model testing the between-years association of dominance with bib size and age in relation to sex. The models were computed using both-sexes dominance hierarchies (Table S8 and S9), between-males dominance hierarchies (Table S10 and S11) and between females dominance hierarchies (Table S12). For each dominance hierarchies, the models were computed both with and without the recommended threshold of ratio of interactions to number of individuals in the group superior to 10 (Sánchez-Tójar et al., 2018) (Table S9 and S11), except for female dominance hierarchies for which not enough colonies matched the criterion. We also tested, using the package “emmeans” (Lenth et al., 2023), the slopes of the relationship between dominance with bib size and the relationship between dominance and age for each years (Table S13; Figure S5).
ix. The influence of year 2013 and 2016 on the relationship between dominance and bib size in between-females dominance hierarchies (Table S14).
x. Timing of data collection for bib size and dominance and proportion of individuals per colony per year observed during dominance video collection (Table S15).

## Supplementary 1: Distribution of the normalised dominance score in the dataset

**Figure S1:**
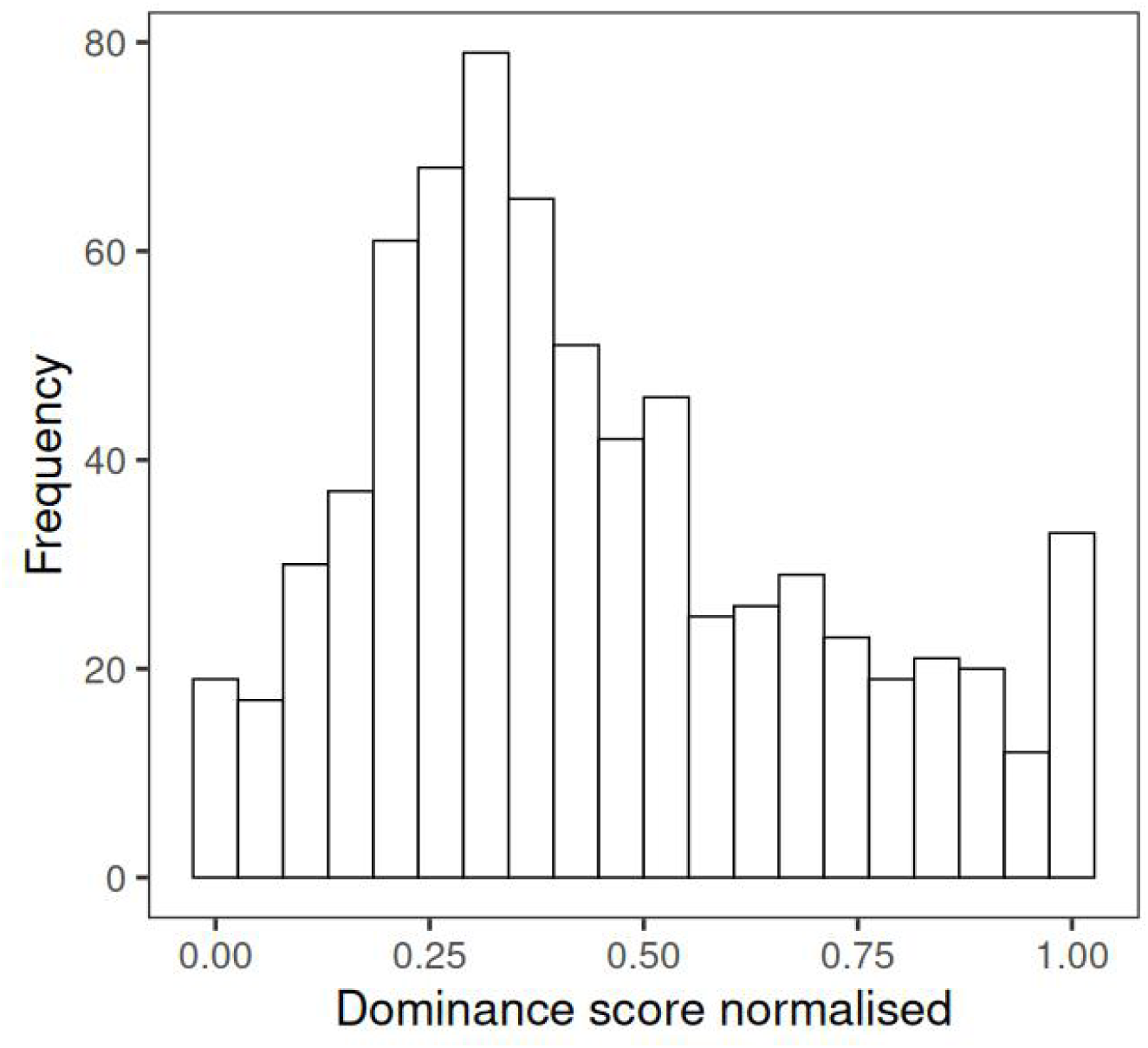
Distribution of the dominance score, normalised by colony by year between zero and one for the 723 individuals present in the dataset. Zero represent the least dominant birds and one the more dominant birds. In the dataset, 13 individuals had a score of exactly zero and 27 had a score of exactly one.

## Supplementary 2: Evaluation of the automatic YOLO method and comparison with previous manual annotations

YOLO is a deep learning model initially specialised in object detection (Redmon et al., 2016), whereas YOLOv8 allows to perform image classification, object segmentation, and pose estimation (Jocher et al., 2022). We used YOLOv8l, a sub-model of YOLOv8, to perform bib segmentation on the birds, due to a good trade-off between number of parameters and processing speed (Jocher et al., 2022; Terven & Cordova-Esparza, 2023).

To train our fine-tuned YOLOv8l model, 1037 pictures were manually annotated using VGG Annotator version 1.0.6 (Dutta & Zisserman, 2019) and we transformed the annotations into the YOLO input format. We separated the dataset into 934 train pictures and 103 validation pictures. Since the pictures were highly standardised, we did not used a test dataset as the validation was enough in these conditions. The model was trained using batches of 14 pictures, an initial learning rate of 0.01 and a final learning rate of 0.001, and data augmentation (flip left/right with a probability of 0.5 on each picture). We used a different image size input from the original YOLOv8 model because we needed accurate segmentation on high quality pictures. Therefore, we defined an image size input of 960×960 pixels.

To evaluate the model we used an Average Precision (AP) metric. The AP computes the number of True Positives divided by the number of True Positives + False Negatives. However, evaluating a segmentation is more complex than a common classification as we need to determine when the segmentation is good enough. For that we use the Intersection over Union (IoU), which is the ratio of the intersection between the true location of the object and the predicted location divided by the union of both the true location and the predicted one. The higher this ratio, the higher is the quality of the detection. An IoU threshold can be set up depending on how accurate the model needs to be. Here, we set up the threshold to 0.9, as we need a very precise overlap between true locations and predicted locations (threshold is usually set to 0.5 when using object detection). At a IoU of 0.9, we obtained an AP of 0.98, which indicates that nearly 100% of the bibs in the 103 validation pictures were at least segmented with 90% of their real location.

The model’s segmentation with YOLO presented similar high repeatability between consecutive pictures of a same individual with the image graphic editor GIMP: YOLO, (mean±SE) 0.90±0.016; CI: [0.866, 0.93]; P<0.001; N=300 versus GIMP manual observer: (mean±SE) 0.92±0.014; CI: [0.885, 0.941]; P<0.001; N=300. The speed to process 30 pictures was also compared, and the deep learning model showed much faster performance than the GIMP method: (mean±SD) 10.2±0.04 seconds to process 30 pictures using the YOLOv8l model (N=10 repetitions) versus 21min07s seconds for the GIMP method (N=1 repetition). Additionally, the accuracy of detection between real bib sizes (VGG annotations), the model’s segmentation, and the previous method (GIMP measures) were compared. Paired T-tests and correlations tests were performed on 82 pictures and the GIMP method seems to be closer to the VGG annotations than the YOLO method, but the YOLO model was slightly more correlated with the VGG annotations (Table S1).

These tests allow us to validate the method using deep learning object segmentation for fine-scale measurement of phenotypic traits. In the dataset used for the analysis, only six years out of seven have the bib size computed using the YOLO method. The GIMP method was used to obtain the bib sizes of the season 2013-2014 due to a loss of the pictures this year and the impossibility to compute the pictures again with the YOLO method. However, bib sizes obtained using the GIMP and the YOLO methods are highly correlated (Table S1), and as bib sizes are normalized within year by colony, results still be comparable.

We also evaluated the reliability of the bib sizes obtained using pictures. We blindly selected 27 individuals for which we computed the repeatability between two to three set of three pictures taken a few dozens of minutes apart. We used the average bib size, extracted with the YOLO method, on the consecutive three pictures per individuals and we compared the replicates, for which we obtained high repeatability (0.838±0.06 (mean±SE); CI: [0.723, 0.93]; P<0.001, N=58).

**Table S1:**
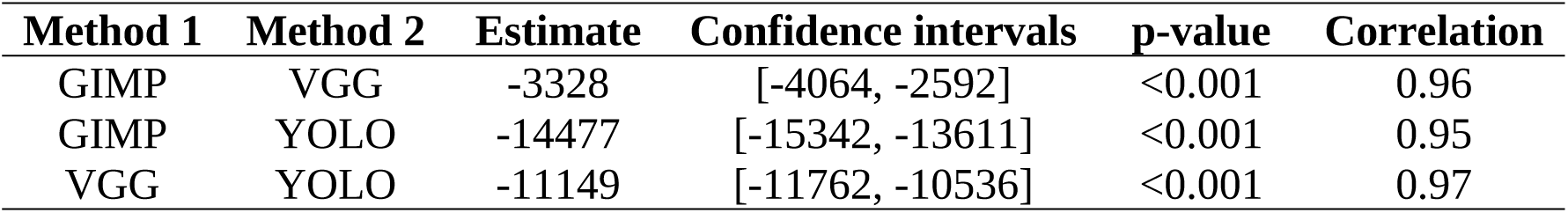
Paired T-test and correlation test between the VGG annotations, the GIMP data processing and the YOLOv8l fine-tuned model.

## Supplementary 3: Evaluation of the link between bib size, age and tarsus length

**Table S2:**
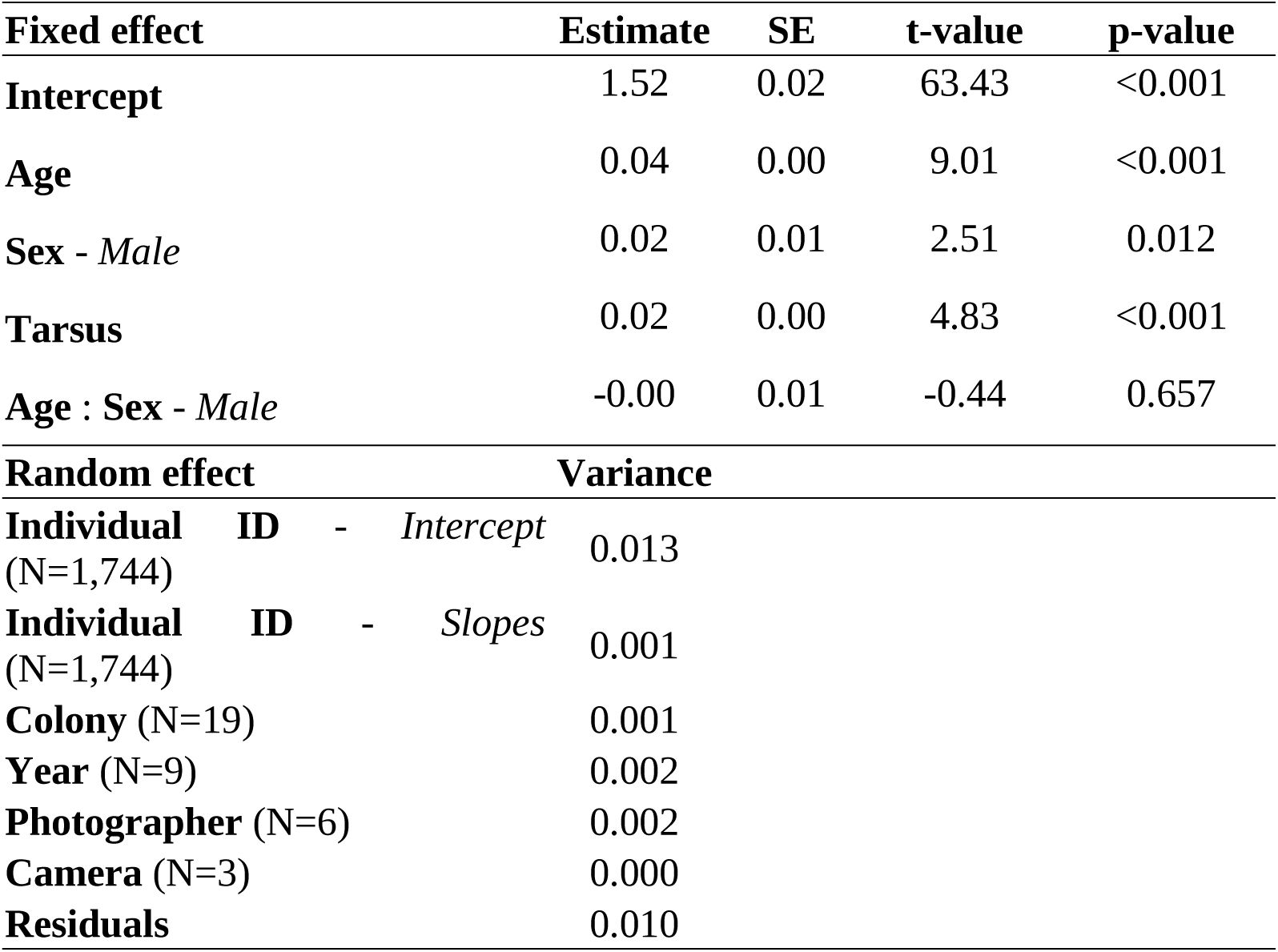
Estimates of the model parameters to test the influence of the size and the age on the bib size without one Photographer that was responsible for variance deviation. Bib size was taken once a year at the end of winter and some individuals were collected multiples years. The model was computedon a reduced dataset without one of the scorers (N=3,373 observations, 814 males and 930 females). The model did meet the residuals normality assumption. Here, ‘Age’ is either exact age for individuals born on our studied colonies or minimum age for individuals first caught as adults.

**Table S3:**
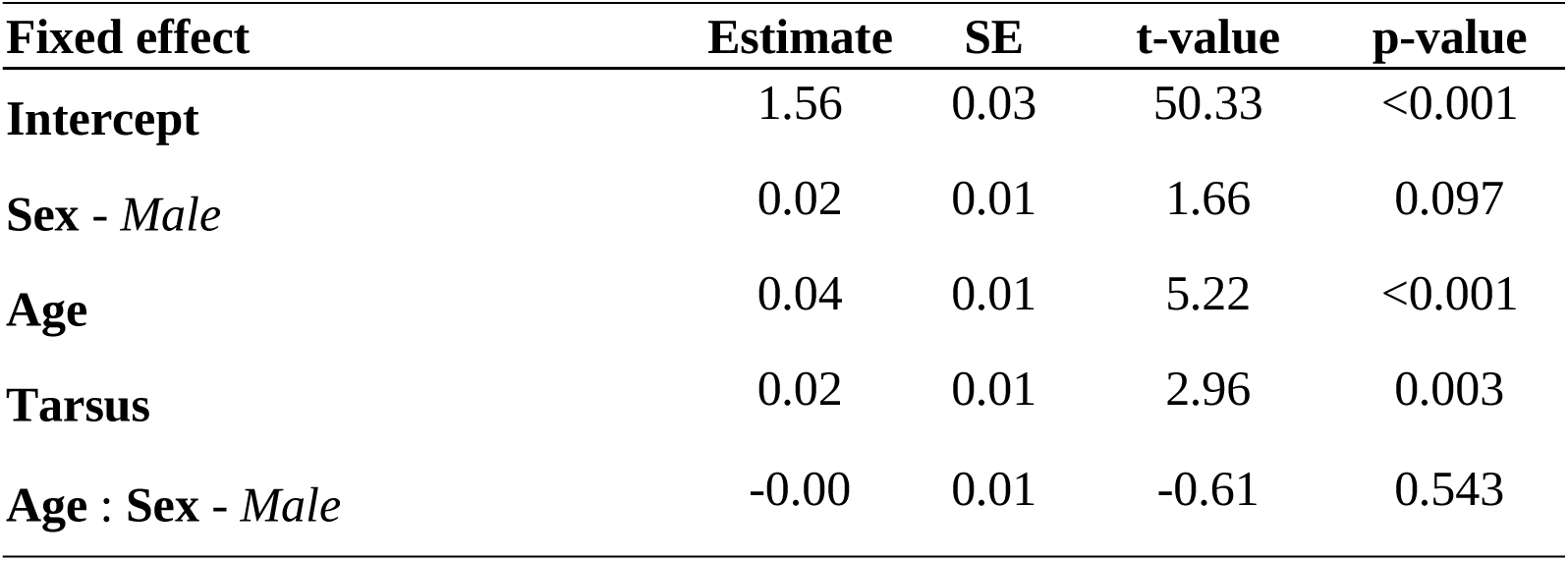
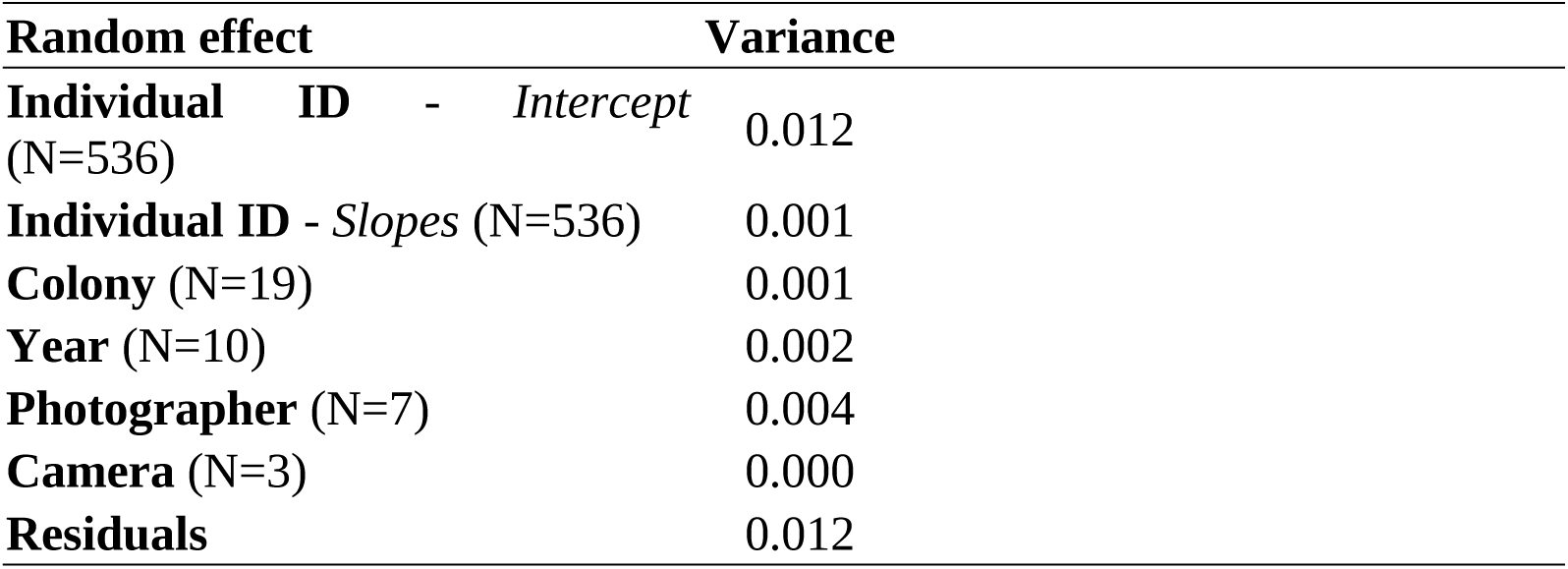
Estimates of the model parameters to test the influence of the size and the age on the bib size with a subset of individual with at least three measures per individuals ((mean ±SD) 3.88±1.15 measures per individuals). The table is the summary of the effects using 2,081 observations (313 males and 223 females). Here, ‘Age’ is either exact age for individuals born on our studied colonies or minimum age for individuals first caught as adults.

## Supplementary 4: Directed Acyclic Graph of the model testing the effect of the bib on dominance

**Figure S2:**
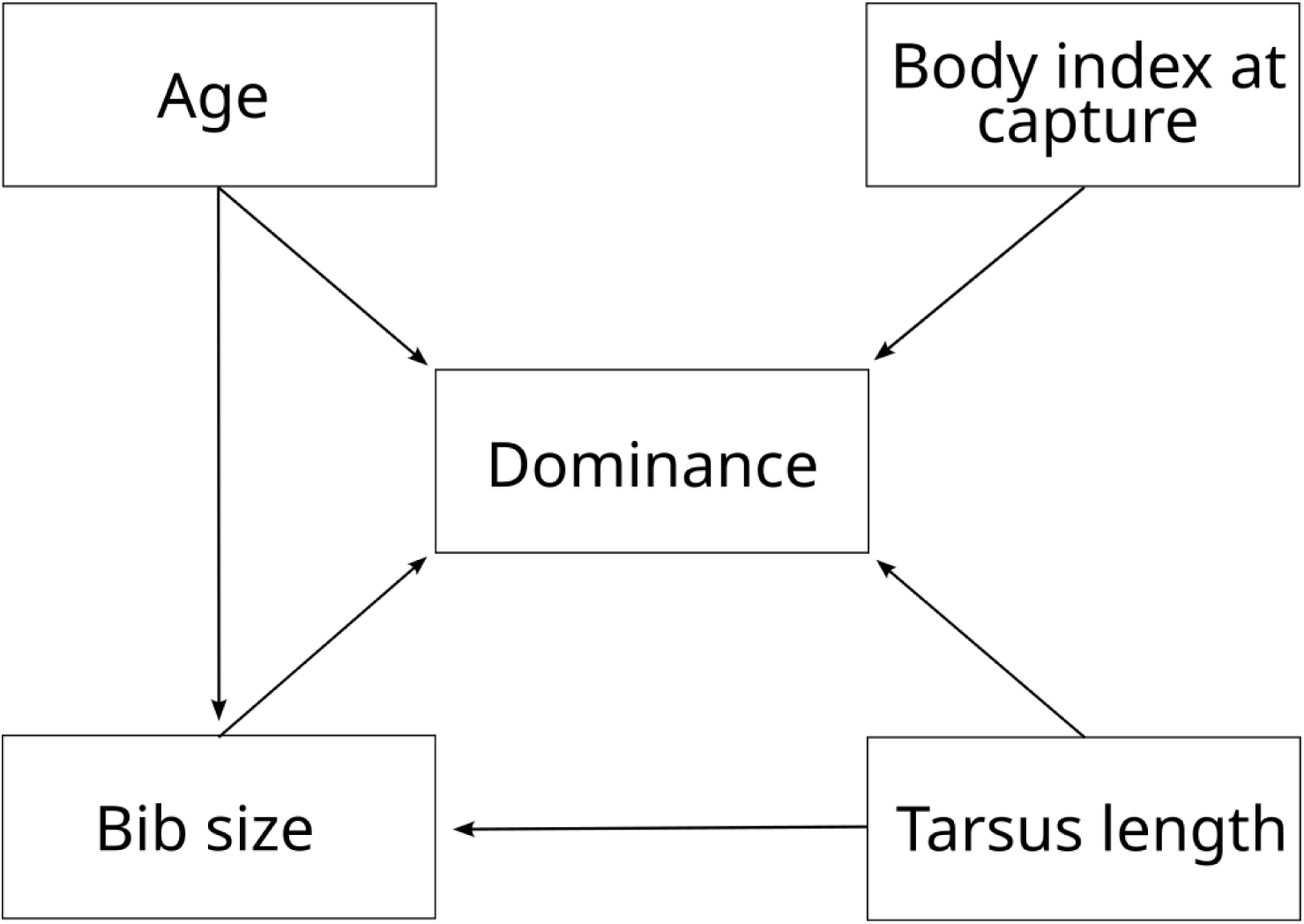
DAG of the model testing the factors impacting dominance. Arrows represent the direction of the effect. In the model structure, interactions between age and sex, and bib size and sex were added as fixed factors. Colony ID, individual identity and year of the data collection were added as random factors.

## Supplementary 5: Association between bib size and tarsus length

**Figure S3:**
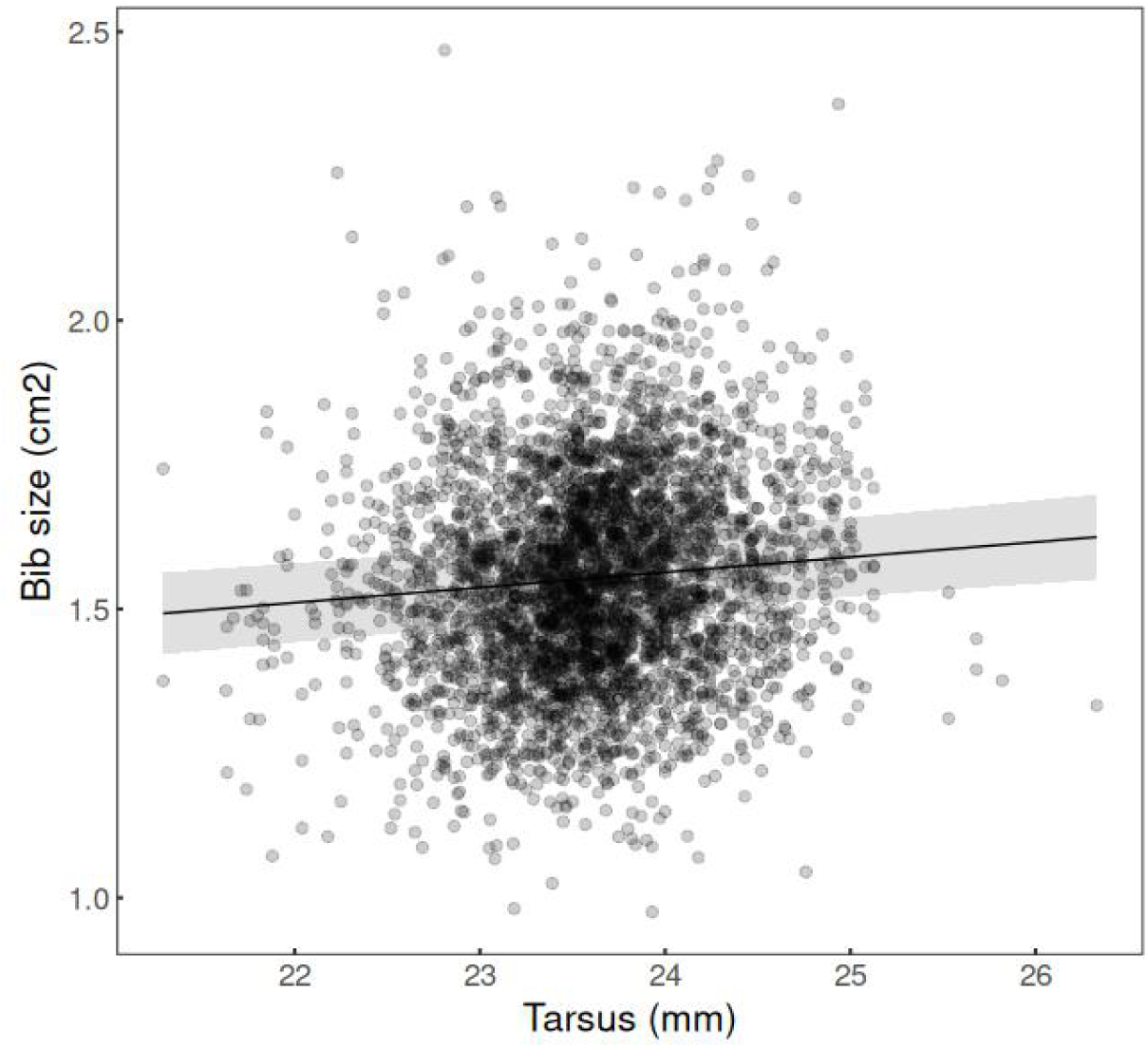
Association between bib size (cm²) and tarsus (mm). Dots correspond to raw data and lines represent the estimated linear fit and its 95% confidence intervals from the mixed model in Table 1.

## Supplementary 6: Justification of the use of Gaussian distribution models to test for the association between dominance and bib size and age

For the model testing the association between dominance and the different factors, we used a Linear Mixed Model which implies Gaussian distribution. Initially, we aimed at a Beta distribution model, adapted to estimate distribution between 0 and 1, and because Beta distribution excludes exact 0 and 1, we added and subtracted 10-10 to respectively 0 and 1. However, the model did not fit well (see analyses of the residuals; Fig. S4). The distribution of the dependent variable is not what determines the link function to be used in the model, and it is only an indication of the model type that we could use but it is not what it determines the link function chosen. It is true that the estimated distribution will not be bounded by 0 and 1 as our data is, but since these data point are relatively rare (13 points with a score of zero and 27 with a score of one), lower and higher estimates will be rare. Analyses of the residuals of the model (Fig. S4) show that the Linear Mixed Model using a Gaussian link fitted better than the Beta distribution model and fitted particularly well with no statistical deviation from normality, no statistical over-dispersion of the residuals and no abnormal presence of outliers.

**Figure S4:**
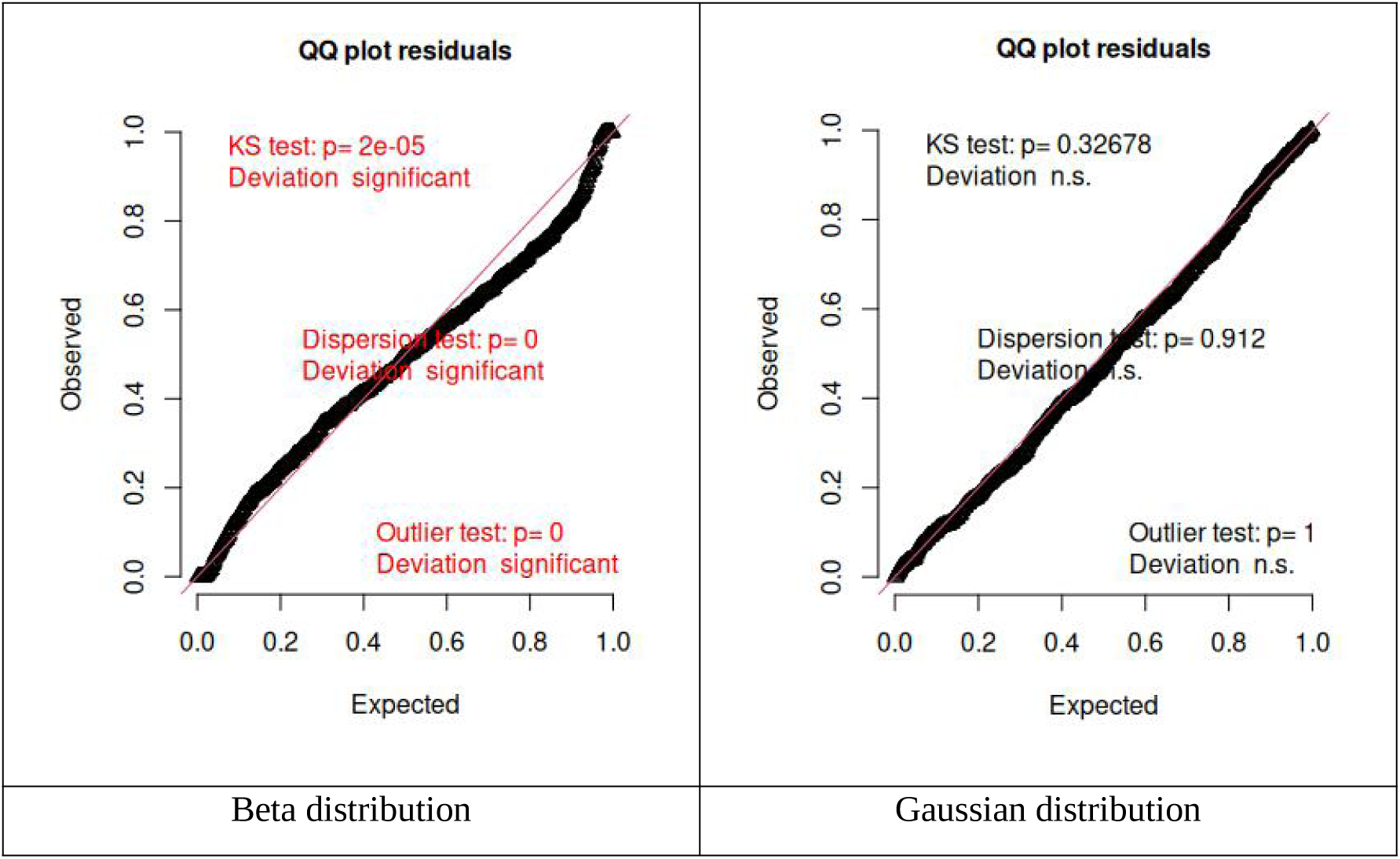
QQ-plot analyses of the residuals of the mixed model testing the association between dominance with bib size and age using the both-sexes dominance hierarchies. The model was initially tested using a Beta distribution model that did not fit well our data (see QQ-plot on the right). The Gaussian distribution model fitted better and fitted well (see QQ-plot on the left). The Gaussian distribution model was therefore kept for the analyses.

## Supplementary 7: Evaluation of the effect of the ratio of dyads per individuals on the overall association between dominance and bib size

Using only the colonies with a ration of number of interactions to individuals over 10, no major differences were observed in the values and directions of the estimates of the significant effects in both-sexes dominance hierarchies and in between-males dominance hierarchies (Table S4 to S6). No model was computed for between-females dominance hierarchies as only one colony one year respected the minimum ratio of interactions to individuals superior to 10. Both the models using both-sexes dominance hierarchies and between-males dominance hierarchies presented an improvement of the models’ assumptions for validation with less heteroscedasticity at the first and third quantile of the residuals when being computed with the colonies with a ration of number of interactions to individuals over 10 (Table S4 to S6). In these models, Age was either exact age for individuals born on our studied colonies or minimum age for individuals first caught as adults.

**Table S4:** Estimates of the model testing the overall influence of bib size, sex and age on dominance in both-sexes dominance hierarchies, without colonies with a ratio inferior to 10 of interactions to the number of individuals. The model was computed with 612 observations. The model’s residuals followed normality, but heteroscedasticity was observed near the residuals’ quantiles 0.25 and 0.75.

| **Fixed effect** | **Estimate** | **SE** | **t-value** | **p-value** |
| --- | --- | --- | --- | --- |
| **Intercept** | 0.26 | 0.04 | 6.87 | <0.001 |
| **Bib size** | 0.01 | 0.01 | 0.80 | 0.423 |
| **Sex - Male** | 0.30 | 0.02 | 19.05 | <0.001 |
| **Age** | 0.00 | 0.01 | 0.38 | 0.702 |
| **Body condition** | 0.01 | 0.01 | 1.01 | 0.311 |
| **Tarsus** | 0.00 | 0.01 | 0.31 | 0.755 |
| **Bib size : Sex - Male** | 0.00 | 0.01 | -0.04 | 0.968 |
| **Age : Sex - Male** | 0.11 | 0.02 | 7.00 | <0.001 |
| **Random effect** | **Variance** |  |  |  |
| **Individual ID (N=377)** | 0.008 |  |  |  |
| **Year (N=7)** | 0.004 |  |  |  |
| **Colony (N=6)** | 0.004 |  |  |  |
| **Residuals** | 0.020 |  |  |  |

**Table S5:** Estimates of the model testing the overall influence of bib size, sex and age on dominance in between-males dominance hierarchies, with colonies with a ratio inferior to 10 of interactions to the number of individuals. The model was computed with 419 observations. The model’s residuals did not follow normality, and heteroscedasticity was observed near the residuals’ quantiles 0.25, 0.5 and 0.75. Association of dominance with bib size and sex is visually represented in Fig. 2.E and 2.F.

| **Fixed effect** | **Estimate** | **SE** | **t-value** | **p-value** |
| --- | --- | --- | --- | --- |
| **Intercept** | 0.49 | 0.04 | 13.08 | <0.001 |
| **Bib size** | -0.02 | 0.01 | -1.69 | 0.092 |
| **Age** | 0.16 | 0.01 | 11.77 | <0.001 |
| **Body condition** | 0.03 | 0.02 | 1.53 | 0.128 |
| **Tarsus** | 0.01 | 0.02 | 0.40 | 0.690 |
| **Random effect** | **Variance** |  |  |  |
| **Individual ID (N=207)** | 0.018 |  |  |  |
| **Year (N=7)** | 0.001 |  |  |  |
| **Colony (N=7)** | 0.006 |  |  |  |
| **Residuals** | 0.034 |  |  |  |

**Table S6:** Estimates of the model testing the overall influence of bib size, sex and age on dominance in between-males dominance hierarchies, without colonies with a ratio inferior to 10 of interactions to the number of individuals. The model was computed with 188 observations. The model’s residuals followed normality, but heteroscedasticity was observed near the residuals’ quantiles 0.25 and 0.75.

| **Fixed effect** | **Estimate** | **SE** | **t-value** | **p-value** |
| --- | --- | --- | --- | --- |
| **Intercept** | 0.45 | 0.03 | 13.67 | <0.001 |
| **Bib size** | -0.01 | 0.02 | -0.47 | 0.640 |
| **Age** | 0.18 | 0.02 | 9.51 | <0.001 |
| **Body condition** | -0.01 | 0.03 | -0.42 | 0.673 |
| **Tarsus** | -0.05 | 0.03 | -1.84 | 0.067 |

| Random effect | Variance |
| --- | --- |
| Individual ID (N=130) | 0.021 |
| Year (N=7) | 0.005 |
| Colony (N=4) | 0.000 |
| Residuals | 0.026 |

**Table S7:** Estimates of the model testing the overall influence of bib size, sex and age on dominance in between-females dominance hierarchies, with colonies with a ratio inferior to 10 of interactions to the number of individuals. The model was computed with 249 observations. The model’s residuals followed normality, and small heteroscedasticity was observed near the residuals’ quantiles 0.5. Association of dominance with bib size and sex is visually represented in Fig. 2.C and 2.D.

| Fixed effect | Estimate | SE | t-value | p-value |
| --- | --- | --- | --- | --- |
| Intercept | 0.50 | 0.03 | 19.63 | <0.001 |
| Bib size | 0.04 | 0.02 | 2.34 | 0.020 |
| Age | -0.01 | 0.02 | -0.35 | 0.725 |
| Body condition | 0.00 | 0.03 | 0.12 | 0.907 |
| Tarsus | -0.01 | 0.03 | -0.53 | 0.598 |

| Random effect | Variance |
| --- | --- |
| Individual ID (N=180) | 0.023 |
| Year (N=7) | 0.001 |
| Colony (N=6) | 0.001 |
| Residuals | 0.048 |

## Supplementary 8: Evaluation of the effect of the ratio of dyads per individuals on the between years association between dominance and bib size

In both-sexes dominance hierarchies, using only the colonies with a ratio of number of interactions to individuals over 10, the sign and estimates of the significant effects did not change (Table S8 and S9), but the interaction between bib size and sex became significant for one season (Table S9). However, (i) the significance of this interactions this year is still in the same direction than the reference year in the model, making the correlation between bib and sex for male being different than the relation between bib and sex for female this year; (ii) the model presented a non-normality of the residuals, potentially affecting the reliability of this additional significant factor. In between-males dominance hierarchies, the signs of the significant effects remained the same between the models using the full dataset and the one using the dataset without the colonies with a ratio of number of interactions to individuals over 10, but the estimates of one of the season levels differed (0.14±0.04 (EST±SE), P=0.002; Table S10; and 0.34±0.09 (EST±SE), P=<0.001; Table S11). No model was computed for the between-females interactions dataset only as only one colony one year respected the minimum ratio of interactions to individuals superior to 10. In these models, Age was either exact age for individuals born on our studied colonies or minimum age for individuals first caught as adults.

**Table S8:**
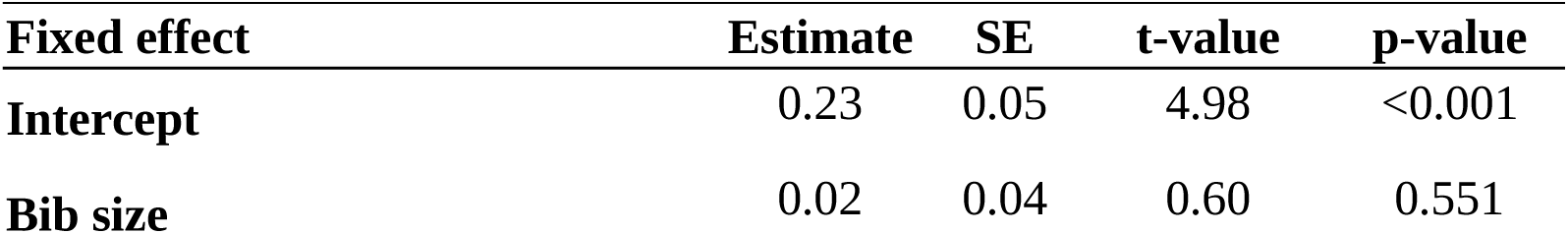
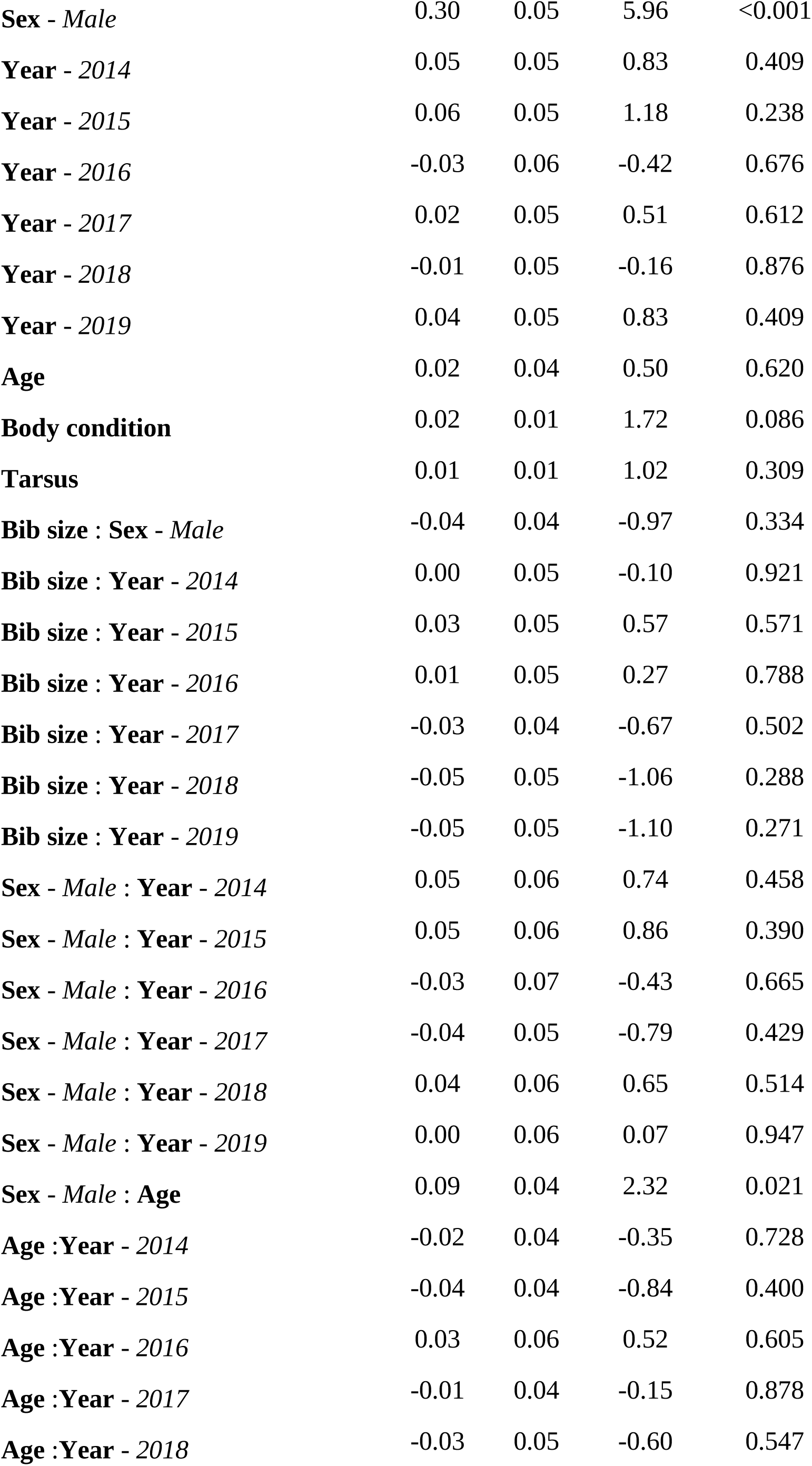
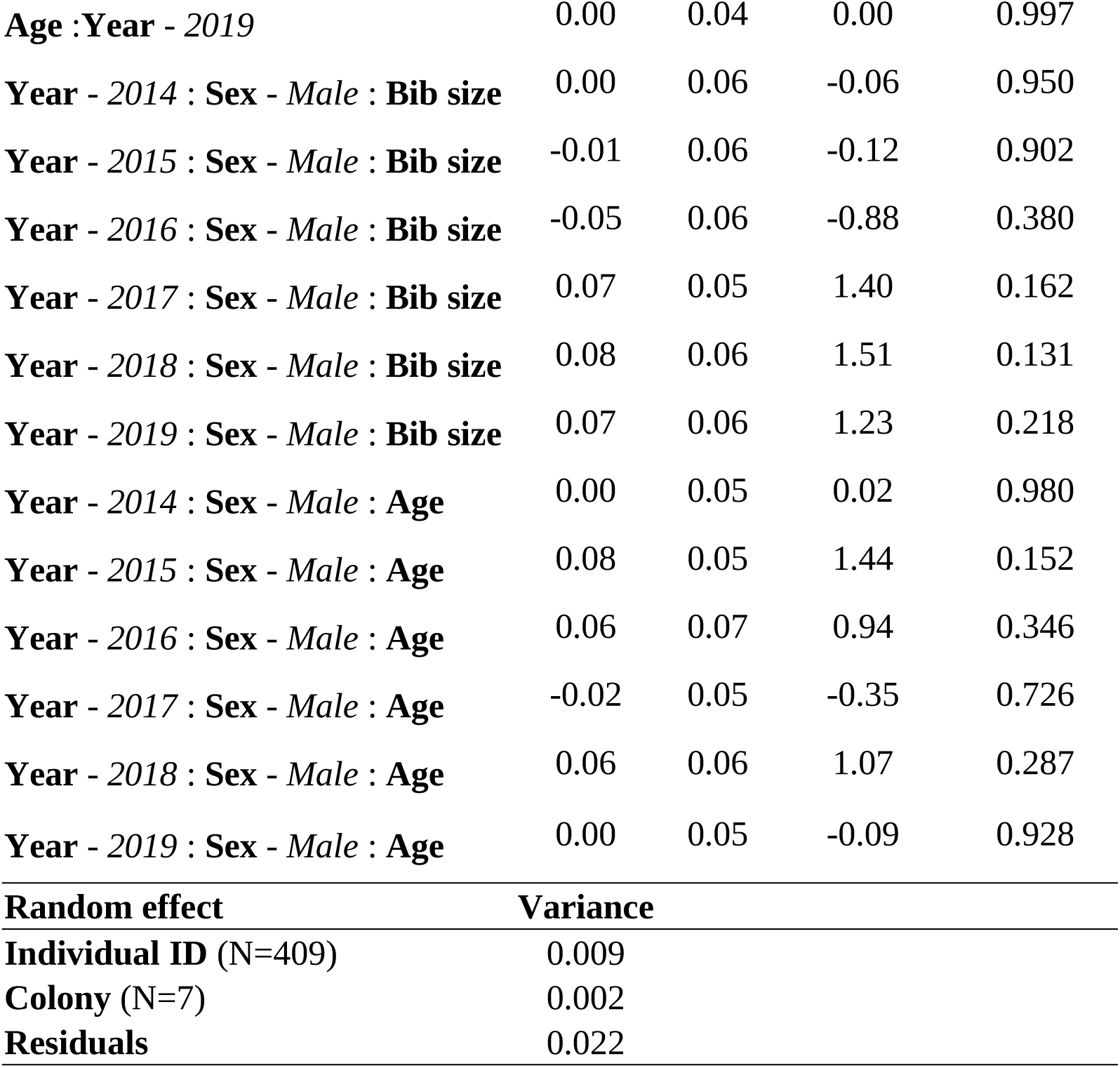
Estimates of the model testing the between years influence of bib size, sex and age on dominance in both-sexes dominance hierarchies, with colonies with a ratio inferior to 10 of interactions to the number of individuals. The model was computed with 723 observations. The model’s residuals followed normality, but heteroscedasticity was observed near the residuals’ quantile 0.75. Association of dominance with bib size and sex is visually represented in Fig. 3.A and 3.B.

**Table S9:** Estimates of the model testing the between years influence of bib size, sex and age on dominance in both-sexes dominance hierarchies, without colonies with a ratio inferior to 10 of interactions to the number of individuals. The model was computed with 612 observations. The model’s residuals did not follow normality, and heteroscedasticity was observed near the residuals’ quantiles 0.25 and 0.75.

| **Fixed effect** | **Estimate** | **SE** | **t-value** | **p-value** |
| --- | --- | --- | --- | --- |
| **Intercept** | 0.24 | 0.05 | 4.73 | <0.001 |
| **Bib size** | 0.01 | 0.04 | 0.25 | 0.799 |
| **Sex - Male** | 0.30 | 0.05 | 6.14 | <0.001 |
| **Year - 2014** | 0.06 | 0.06 | 0.99 | 0.321 |
| **Year - 2015** | 0.05 | 0.05 | 1.04 | 0.297 |
| **Year - 2016** | -0.04 | 0.07 | -0.57 | 0.572 |
| **Year - 2017** | 0.01 | 0.05 | 0.32 | 0.750 |
| **Year - 2018** | -0.08 | 0.05 | -1.48 | 0.139 |
| **Year - 2019** | 0.11 | 0.06 | 1.95 | 0.052 |
| **Age** | 0.02 | 0.03 | 0.68 | 0.495 |
| **Body condition** | 0.01 | 0.01 | 0.58 | 0.563 |
| **Tarsus** | 0.00 | 0.01 | -0.02 | 0.982 |
| **Bib size : Sex - Male** | -0.03 | 0.04 | -0.64 | 0.521 |
| **Bib size : Year - 2014** | 0.02 | 0.05 | 0.35 | 0.727 |
| **Bib size : Year - 2015** | 0.04 | 0.05 | 0.78 | 0.434 |
| **Bib size : Year - 2016** | 0.02 | 0.06 | 0.37 | 0.709 |
| **Bib size : Year - 2017** | -0.01 | 0.04 | -0.21 | 0.833 |
| **Bib size : Year - 2018** | 0.00 | 0.05 | 0.01 | 0.990 |
| **Bib size : Year - 2019** | -0.03 | 0.05 | -0.59 | 0.556 |
| **Sex - Male : Year - 2014** | 0.03 | 0.07 | 0.50 | 0.614 |
| **Sex - Male : Year - 2015** | 0.05 | 0.06 | 0.87 | 0.384 |
| **Sex - Male : Year - 2016** | -0.03 | 0.08 | -0.37 | 0.711 |
| **Sex - Male : Year - 2017** | -0.04 | 0.05 | -0.83 | 0.408 |
| **Sex - Male : Year - 2018** | 0.03 | 0.06 | 0.47 | 0.640 |
| **Sex - Male : Year - 2019** | -0.02 | 0.07 | -0.30 | 0.765 |
| **Sex - Male : Age** | 0.08 | 0.04 | 2.14 | 0.033 |
| **Age : Year - 2014** | -0.02 | 0.04 | -0.36 | 0.722 |
| **Age : Year - 2015** | -0.05 | 0.04 | -1.07 | 0.285 |
| **Age : Year - 2016** | -0.05 | 0.07 | -0.75 | 0.451 |
| **Age : Year - 2017** | -0.01 | 0.04 | -0.13 | 0.897 |
| **Age : Year - 2018** | -0.01 | 0.05 | -0.22 | 0.826 |
| **Age : Year - 2019** | -0.03 | 0.04 | -0.74 | 0.462 |
| **Year - 2014 : Sex - Male : Bib size** | 0.02 | 0.06 | 0.26 | 0.799 |
| **Year - 2015 : Sex - Male : Bib size** | -0.01 | 0.06 | -0.24 | 0.808 |
| **Year - 2016 : Sex - Male : Bib size** | -0.06 | 0.07 | -0.9 | 0.371 |
| **Year - 2017 : Sex - Male : Bib size** | 0.06 | 0.05 | 1.09 | 0.277 |
| **Year - 2018 : Sex - Male : Bib size** | 0.05 | 0.06 | 0.80 | 0.424 |
| **Year - 2019 : Sex - Male : Bib size** | 0.05 | 0.06 | 0.95 | 0.341 |
| **Year - 2014 : Sex - Male : Age** | -0.02 | 0.05 | -0.37 | 0.712 |
| **Year - 2015 : Sex - Male : Age** | 0.09 | 0.05 | 1.76 | 0.078 |
| **Year - 2016 : Sex - Male : Age** | 0.18 | 0.09 | 1.99 | 0.047 |
| **Year - 2017 : Sex - Male : Age** | 0.00 | 0.05 | -0.10 | 0.918 |
| **Year - 2018 : Sex - Male : Age** | 0.06 | 0.06 | 0.97 | 0.334 |
| **Year - 2019 : Sex - Male : Age** | 0.03 | 0.06 | 0.50 | 0.615 |

| **Random effect** | **Variance** |
| --- | --- |
| **Individual ID (N=377)** | 0.008 |
| **Colony (N=6)** | 0.004 |
| **Residuals** | 0.020 |

**Table S10:** Estimates of the model testing the between years influence of bib size, sex and age on dominance in between-males dominance hierarchies, with colonies with a ratio inferior to 10 of interactions to the number of individuals. The model was computed with 419 observations. The model’s residuals followed normality, but heteroscedasticity was observed near the residuals’ quantiles 0.25, 0.5 and 0.75. Association of dominance with bib size and sex is visually represented in Fig. 3.E and 3.F.

| **Fixed effect** | **Estimate** | **SE** | **t-value** | **p-value** |
| --- | --- | --- | --- | --- |
| **Intercept** | 0.42 | 0.05 | 8.89 | <0.001 |
| **Bib size** | -0.02 | 0.03 | -0.64 | 0.519 |
| **Year - 2014** | 0.14 | 0.04 | 3.24 | 0.001 |
| **Year - 2015** | 0.12 | 0.04 | 2.73 | 0.007 |
| **Year - 2016** | 0.06 | 0.05 | 1.32 | 0.188 |
| **Year - 2017** | 0.03 | 0.04 | 0.80 | 0.427 |
| **Year - 2018** | 0.08 | 0.05 | 1.61 | 0.108 |
| **Year - 2019** | 0.07 | 0.05 | 1.43 | 0.154 |
| **Age** | 0.16 | 0.02 | 6.39 | <0.001 |
| **Body condition** | 0.03 | 0.02 | 1.43 | 0.155 |
| **Tarsus** | 0.01 | 0.02 | 0.45 | 0.656 |
| **Bib size : Year - 2014** | -0.04 | 0.04 | -1.02 | 0.310 |
| **Bib size : Year - 2015** | 0.01 | 0.04 | 0.15 | 0.879 |
| **Bib size : Year - 2016** | -0.03 | 0.04 | -0.7 | 0.487 |
| **Bib size : Year - 2017** | 0.02 | 0.04 | 0.51 | 0.609 |
| **Bib size : Year - 2018** | 0.02 | 0.04 | 0.39 | 0.695 |
| **Bib size : Year - 2019** | 0.01 | 0.04 | 0.23 | 0.816 |
| **Age :Year - 2014** | -0.01 | 0.04 | -0.31 | 0.760 |
| **Age :Year - 2015** | 0.03 | 0.04 | 0.79 | 0.428 |
| **Age :Year - 2016** | 0.05 | 0.05 | 0.98 | 0.326 |
| **Age :Year - 2017** | -0.02 | 0.03 | -0.66 | 0.510 |
| **Age :Year - 2018** | 0.00 | 0.04 | -0.08 | 0.937 |
| **Age :Year - 2019** | -0.03 | 0.04 | -0.62 | 0.538 |
| **Random effect** | **Variance** |  |  |  |
| **Individual ID (N=207)** | 0.018 |  |  |  |
| **Colony (N=7)** | 0.006 |  |  |  |
| **Residuals** | 0.035 |  |  |  |

**Table S11:** Estimates of the model testing the between years influence of bib size, sex and age on dominance in between-males dominance hierarchies, without colonies with a ratio inferior to 10 of interactions to the number of individuals. The model was computed with 188 observations. The model’s residuals followed normality, but heteroscedasticity was observed near the residuals’ quantiles 0.25, 0.5 and 0.75.

| **Fixed effect** | **Estimate** | **SE** | **t-value** | **p-value** |
| --- | --- | --- | --- | --- |
| **Intercept** | 0.34 | 0.06 | 5.34 | <0.001 |
| **Bib size** | -0.03 | 0.06 | -0.47 | 0.639 |
| **Year - 2014** | 0.34 | 0.09 | 3.73 | <0.001 |
| **Year - 2015** | 0.14 | 0.07 | 2.15 | 0.033 |
| **Year - 2016** | 0.10 | 0.07 | 1.34 | 0.181 |
| **Year - 2017** | 0.12 | 0.07 | 1.68 | 0.094 |
| **Year - 2018** | 0.04 | 0.08 | 0.47 | 0.635 |
| **Year - 2019** | 0.10 | 0.08 | 1.25 | 0.212 |
| **Age** | 0.13 | 0.05 | 2.88 | 0.005 |
| **Body condition** | -0.02 | 0.03 | -0.59 | 0.558 |
| **Tarsus** | -0.05 | 0.03 | -1.76 | 0.080 |
| **Bib size : Year - 2014** | -0.10 | 0.08 | -1.22 | 0.225 |
| **Bib size : Year - 2015** | 0.01 | 0.06 | 0.21 | 0.832 |
| **Bib size : Year - 2016** | -0.02 | 0.08 | -0.29 | 0.771 |
| **Bib size : Year - 2017** | 0.05 | 0.07 | 0.81 | 0.417 |
| **Bib size : Year - 2018** | 0.09 | 0.07 | 1.27 | 0.205 |
| **Bib size : Year - 2019** | 0.03 | 0.07 | 0.46 | 0.646 |
| **Age :Year - 2014** | 0.09 | 0.08 | 1.13 | 0.261 |
| **Age :Year - 2015** | 0.09 | 0.06 | 1.63 | 0.106 |
| **Age :Year - 2016** | 0.06 | 0.09 | 0.70 | 0.486 |
| **Age :Year - 2017** | 0.08 | 0.06 | 1.27 | 0.207 |
| **Age :Year - 2018** | 0.11 | 0.06 | 1.97 | 0.051 |
| **Age :Year - 2019** | 0.00 | 0.06 | 0.00 | 0.999 |
| **Random effect** | **Variance** |  |  |  |
| **Individual ID (N=130)** | 0.024 |  |  |  |
| **Colony (N=4)** | 0.000 |  |  |  |
| **Residuals** | 0.022 |  |  |  |

**Table S12:** Estimates of the model testing the between years influence of bib size, sex and age on dominance in between-females dominance hierarchies, with colonies with a ratio inferior to 10 of interactions to the number of individuals. The model was computed with 249 observations. The model’s residuals followed normality, but heteroscedasticity was observed near the residuals’ quantile 0.75. Association of dominance with bib size and sex is visually represented in Fig. 3.C and 3.D.

| **Fixed effect** | **Estimate** | **SE** | **t-value** | **p-value** |
| --- | --- | --- | --- | --- |
| **Intercept** | 0.47 | 0.06 | 7.25 | <0.001 |
| **Bib size** | 0.14 | 0.06 | 2.36 | 0.019 |
| **Year - 2014** | -0.05 | 0.09 | -0.53 | 0.593 |
| **Year - 2015** | -0.01 | 0.07 | -0.20 | 0.843 |
| **Year - 2016** | 0.03 | 0.10 | 0.33 | 0.739 |
| **Year - 2017** | 0.07 | 0.07 | 1.07 | 0.287 |
| **Year - 2018** | 0.04 | 0.08 | 0.45 | 0.650 |
| **Year - 2019** | 0.03 | 0.08 | 0.43 | 0.665 |
| **Age** | 0.00 | 0.05 | -0.05 | 0.958 |
| **Body condition** | -0.01 | 0.03 | -0.27 | 0.787 |
| **Tarsus** | -0.02 | 0.03 | -0.83 | 0.405 |
| **Bib size : Year - 2014** | -0.07 | 0.08 | -0.83 | 0.407 |
| **Bib size : Year - 2015** | -0.08 | 0.07 | -1.08 | 0.280 |
| **Bib size : Year - 2016** | 0.04 | 0.10 | 0.39 | 0.696 |
| **Bib size : Year - 2017** | -0.10 | 0.07 | -1.50 | 0.135 |
| **Bib size : Year - 2018** | -0.13 | 0.08 | -1.67 | 0.096 |
| **Bib size : Year - 2019** | -0.13 | 0.07 | -1.89 | 0.060 |
| **Age :Year - 2014** | 0.09 | 0.06 | 1.47 | 0.142 |
| **Age :Year - 2015** | -0.03 | 0.06 | -0.48 | 0.631 |
| **Age :Year - 2016** | 0.11 | 0.10 | 1.05 | 0.297 |
| **Age :Year - 2017** | -0.07 | 0.06 | -1.21 | 0.228 |
| **Age :Year - 2018** | 0.11 | 0.09 | 1.13 | 0.260 |
| **Age :Year - 2019** | 0.00 | 0.06 | -0.07 | 0.946 |
| **Random effect** | **Variance** |  |  |  |
| **Individual ID (N=180)** | 0.025 |  |  |  |
| **Colony (N=6)** | 0.002 |  |  |  |
| **Residuals** | 0.046 |  |  |  |

**Table S13:**
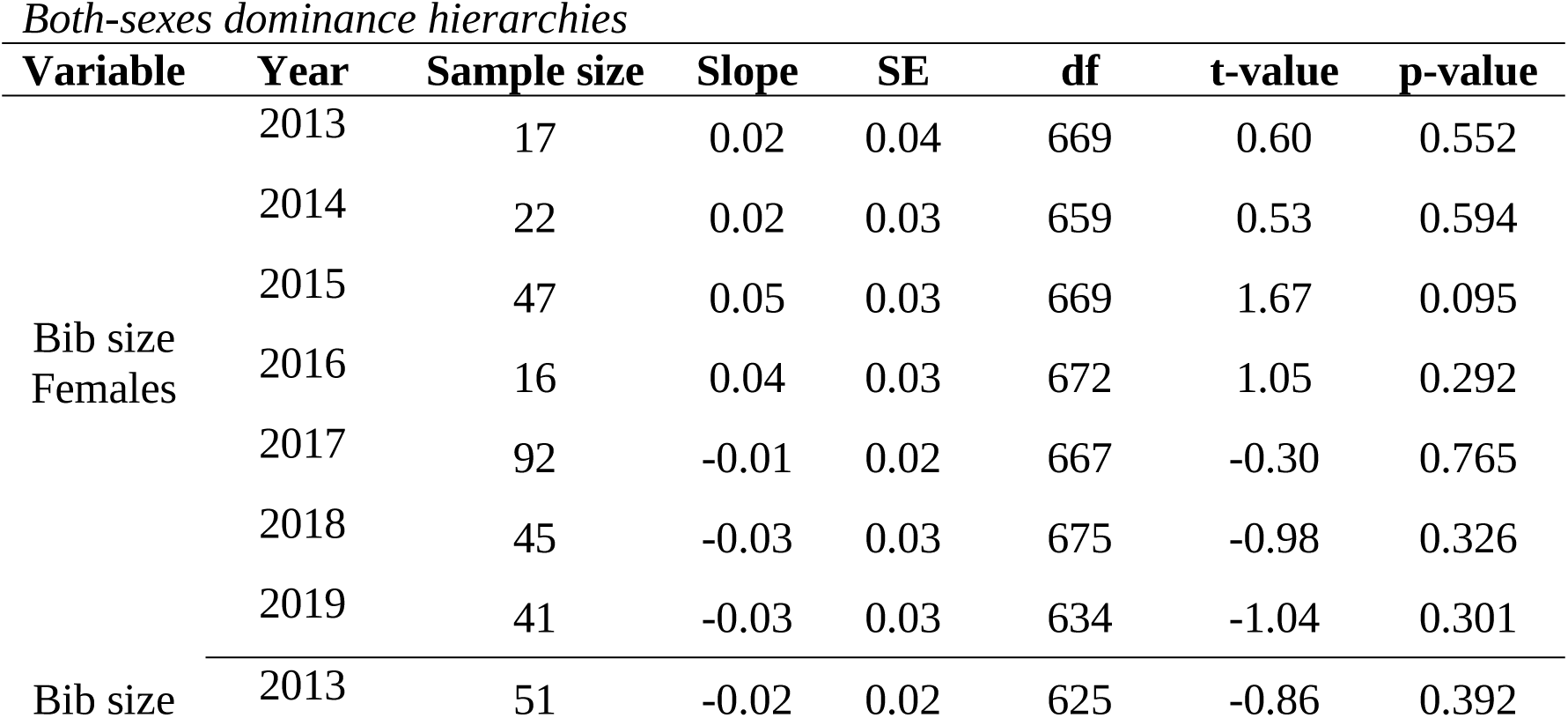
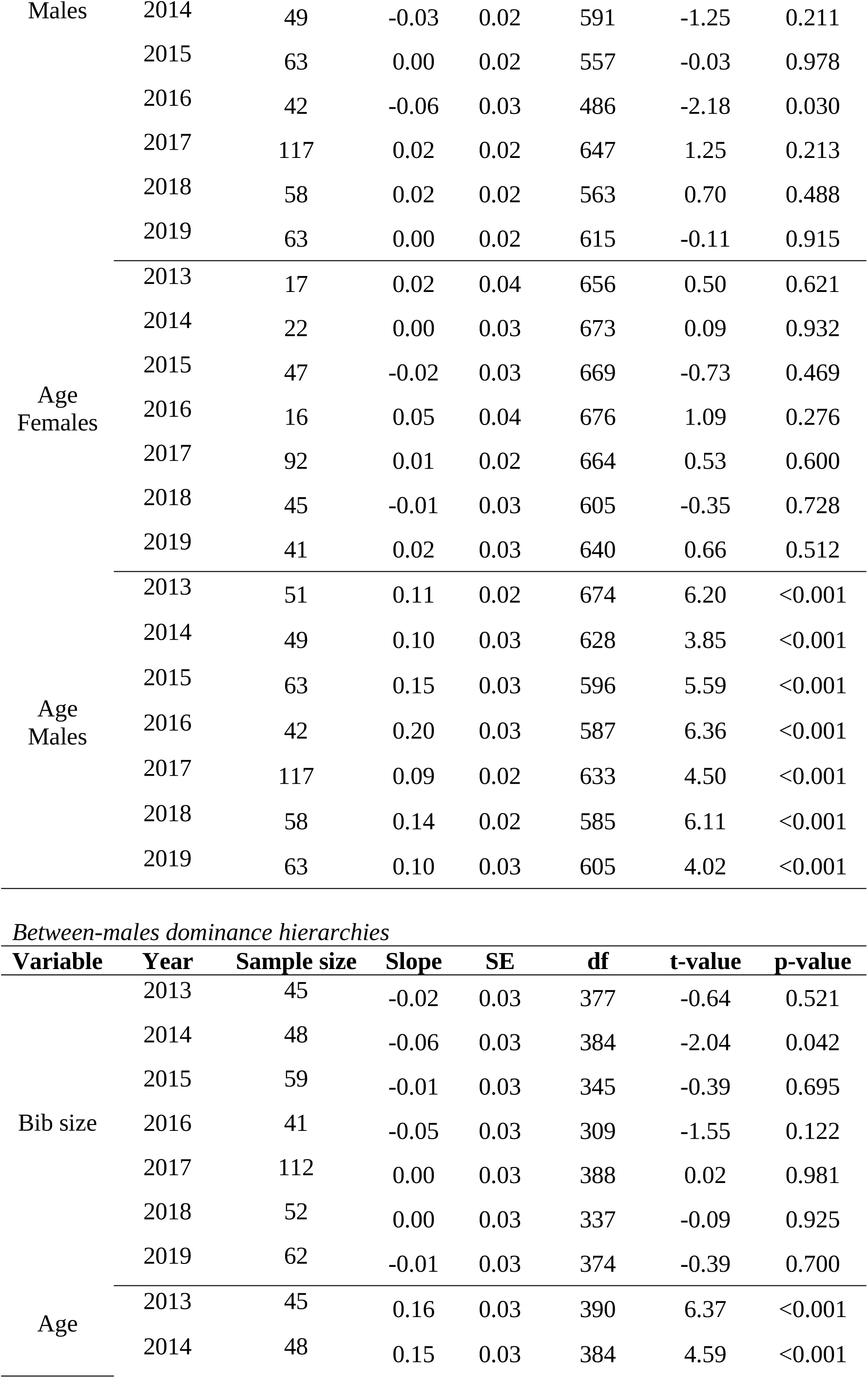
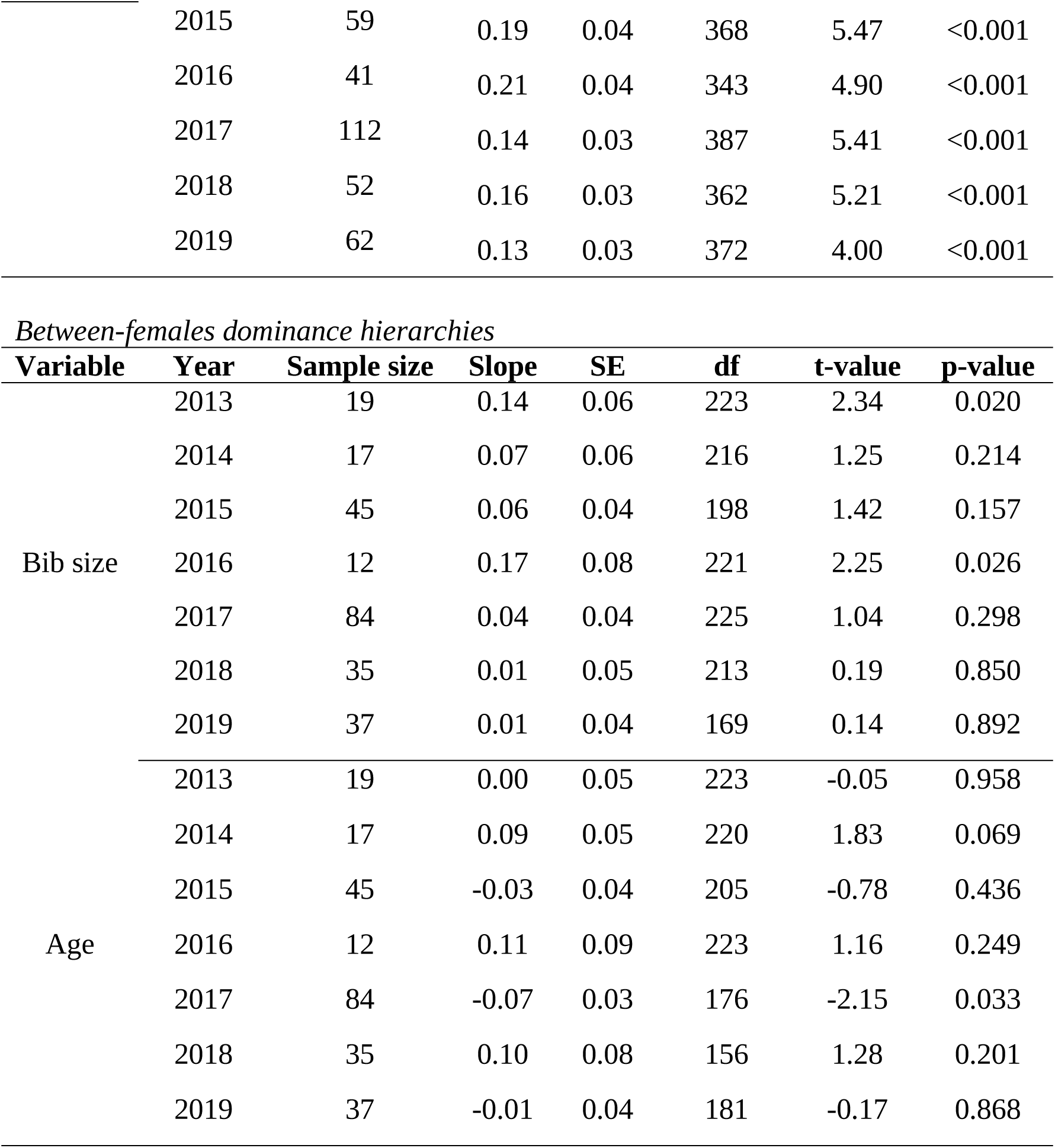
Post-hoc tests analysing bib size’s and age’s slopes of the between-years variation GLMM for both-sexes dominance hierarchies, between-males dominance hierarchies, and between-females dominance hierarchies. Sample size corresponds to the number of individuals present in the dataset for a specific year. The tests were performed with the package “emmeans”.

**Figure S5:**
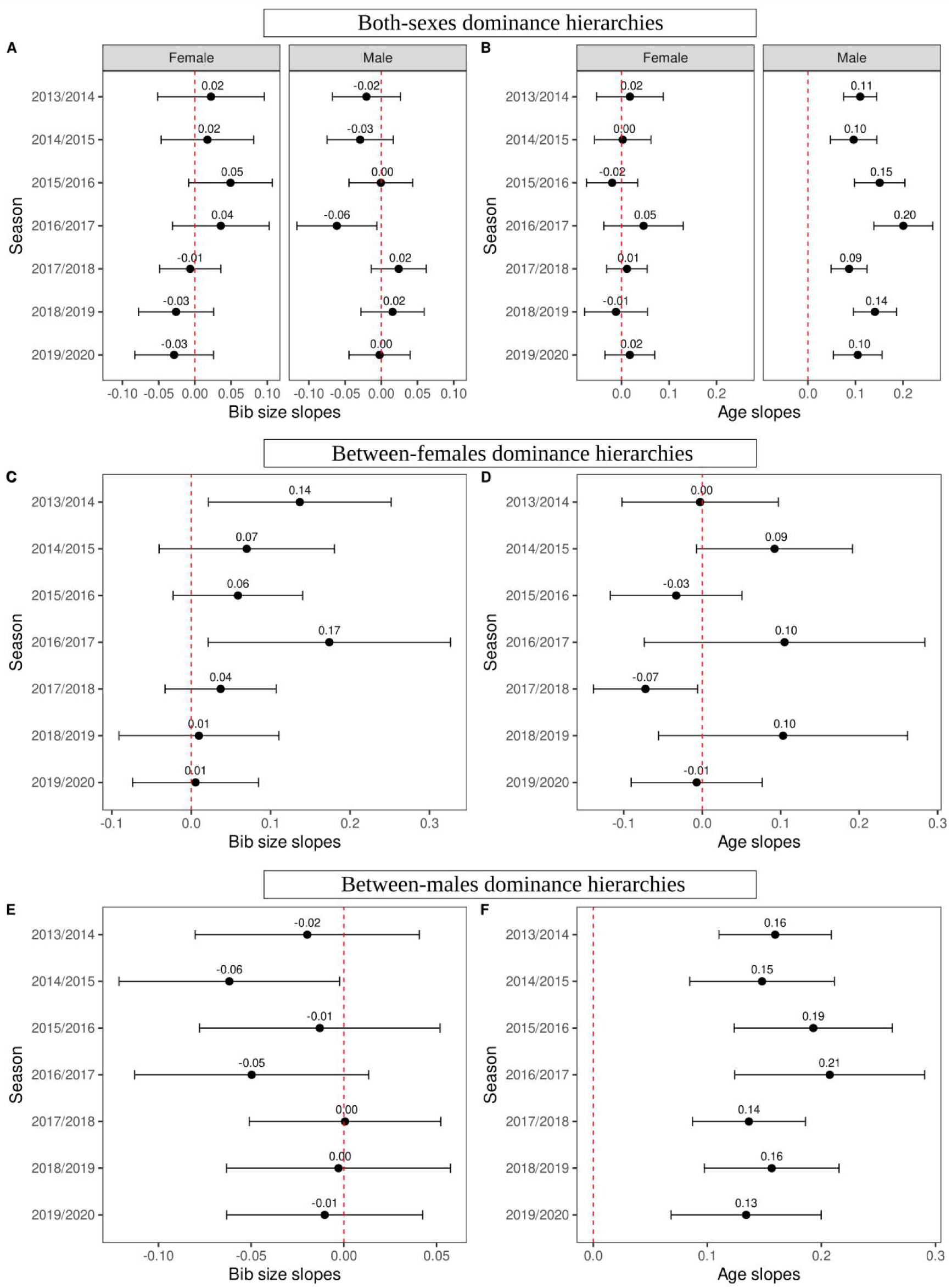
Slopes of the association between: (A,C,E) dominance and normalized bib size; and (B,D,F) dominance and age. Results were obtained from the models using both-sexes dominance hierarchies (A,B), between-female dominance hierarchies (C,D) and between-male dominance hierarchies (E,F). Dots represent the estimated means of the relationships and bar are the confidence intervals.

## Supplementary 9: Influence of year 2013 and 2016 on the relationship between dominance and bib size in between-females dominance hierarchies

**Table S14:** Estimates of the model testing the overall influence of bib size, sex and age on dominance in between-females dominance hierarchies, with colonies with a ratio inferior to 10 of interactions to the number of individuals but without year 2013 and 2016. These two seasons were removed to test their influence on the relationship between dominance and bib size found in Table S7. The model was computed with 219 observations. The model’s residuals followed normality, and small heteroscedasticity was observed near the residuals’ quantiles 0.75.

| Fixed effect | Estimate | SE | t-value | p-value |
| --- | --- | --- | --- | --- |
| Intercept | 0.50 | 0.03 | 17.22 | <0.001 |
| Bib size | 0.02 | 0.02 | 1.06 | 0.290 |
| Age | -0.01 | 0.02 | -0.38 | 0.704 |
| Body condition | 0.01 | 0.03 | 0.46 | 0.646 |
| Tarsus | -0.02 | 0.03 | -0.67 | 0.503 |
| Random effect | Variance |  |  |  |
| Individual ID (N=163) | 0.019 |  |  |  |
| Year (N=5) | 0.001 |  |  |  |
| Colony (N=6) | 0.001 |  |  |  |
| Residuals | 0.047 |  |  |  |

## Supplementary 10: Timing of data collection for bib size and dominance and proportion of individuals per colony per year observed during dominance video collection

**Table S15:** Timing of data collection for bib size and dominance and proportion of individuals per colony per year observed during dominance video collection. Date for bib size data collection are given as month.day. Dominance data collection timing are given as the number of days after bib size collection. The total number of individuals used to compute the proportion of the colony observed is the total number of birds capture at a colony a year during the day of bib size collection. Proportion higher to 1 can happen when more individuals are observed on the videos than caught before (due to dispersal for instance). Stars corresponds to colonies where dominance was sampled more than 40 days after bib size collection date. In total, (mean±SD) 0.85%±0.23% were observed on average per colony per year, and the dominance data were collected on average on a period of (mean±SD) 9±9 days.

| **Season** | **Colony** | **Bib size date** | **Days after bib size collection dominance was collected** | **Number of days recording** | **Proportion of the colony observed** |
| --- | --- | --- | --- | --- | --- |
| 2013 | 11 | 08.31 | 4 to 22 | 16 | 0.80 |
|  | 20* | 09.04 | 59 to 60 and 81 to 92 | 14 | 0.74 |
|  | 21* | 09.09 | 48 to 59 | 12 | 1.00 |
|  | 27 | 09.10 | 23 to 35 | 10 | 0.82 |
|  | 71 | 09.07 | 18 to 30 | 11 | 1.55 |
|  | 81 | 09.13 | 5 and 27 to 40 | 13 | 0.43 |
| 2014 | 11 | 08.25 | 12 to 19 | 5 | 0.66 |
|  | 20 | 08.28 | 15 to 18 | 4 | 0.76 |
|  | 21 | 09.02 | 7 to 20 | 8 | 1.14 |
|  | 27 | 08.30 | 18 to 21 | 4 | 0.76 |
|  | 71 | 09.01 | 5 to 12 | 5 | 0.81 |
|  | 81 | 08.23 | 15 to 34 | 10 | 0.63 |
| 2015 | 11 | 09.04 | -7 to 5 | 9 | 1.16 |
|  | 20 | 08.31 | -1 to 10 | 11 | 0.90 |
|  | 21 | 08.27 | 1 to 13 | 11 | 1.25 |
|  | 27 | 08.29 | 0 to 12 | 10 | 1.07 |
|  | 71 | 09.03 | -2 to 7 | 9 | 0.98 |
| 2016 | 11 | 08.30 | 20 to 25 | 4 | 0.49 |
|  | 20 | 08.25 | 15 to 24 | 4 | 0.67 |
|  | 27 | 08.31 | -1 to 20 | 7 | 0.78 |
|  | 43 | 08.22 | 10 to 35 | 6 | 0.62 |
| 2017 | 11 | 08.25 | 13 to 19 | 5 | 1.00 |
|  | 20 | 08.29 | 9 | 1 | 0.96 |
|  | 27 | 08.30 | 9 | 1 | 0.94 |
|  | 43 | 08.23 | 10 to 12 | 3 | 0.93 |
|  | 71 | 08.26 | 12 | 1 | 0.93 |
| 2018 | 11* | 09.04 | 102 | 1 | 0.54 |
|  | 20* | 08.28 | 108 | 1 | 0.43 |
|  | 27* | 08.23 | 108 to 115 | 4 | 0.78 |
|  | 43* | 08.26 | 105 to 106 | 2 | 0.73 |
|  | 71* | 09.05 | 96 to 103 | 4 | 0.86 |
| 2019 | 11 | 09.08 | 10 to 12 | 3 | 0.98 |
|  | 20 | 08.31 | 18 to 20 | 3 | 0.96 |
|  | 27 | 09.06 | 12 to 14 | 3 | 0.81 |
|  | 43 | 09.02 | 16 to 22 | 3 | 0.70 |
|  | 71 | 09.10 | 9 to 11 | 2 | 0.88 |

